# Single-cell gene regulatory network analysis reveals new melanoma cell states and transition trajectories during phenotype switching

**DOI:** 10.1101/715995

**Authors:** Jasper Wouters, Zeynep Kalender-Atak, Liesbeth Minnoye, Katina I. Spanier, Maxime De Waegeneer, Carmen Bravo González-Blas, David Mauduit, Kristofer Davie, Gert Hulselmans, Ahmad Najem, Michael Dewaele, Florian Rambow, Samira Makhzami, Valerie Christiaens, Frederik Ceyssens, Ghanem Ghanem, Jean-Christophe Marine, Suresh Poovathingal, Stein Aerts

## Abstract

Melanoma is notorious for its cellular heterogeneity, which is at least partly due to its ability to transition between alternate cell states. Similarly to EMT, melanoma cells with a melanocytic phenotype can switch to a mesenchymal-like phenotype. However, scattered emerging evidence indicates that additional, intermediate state(s) may exist. In order to search for such new melanoma states and decipher their underlying gene regulatory network (GRN), we extensively studied ten patient-derived melanoma cultures by single-cell RNA-seq of >39,000 cells. Although each culture exhibited a unique transcriptome, we identified shared gene regulatory networks that underlie the extreme melanocytic and mesenchymal cell states, as well as one (stable) intermediate state. The intermediate state was corroborated by a distinct open chromatin landscape and governed by the transcription factors EGR3, NFATC2, and RXRG. Single-cell migration assays established that this “transition” state exhibits an intermediate migratory phenotype. Through a dense time-series sampling of single cells and dynamic GRN inference, we unraveled the sequential and recurrent arrangement of transcriptional programs at play during phenotype switching that ultimately lead to the mesenchymal cell state. We provide the scRNA-Seq data with 39,263 melanoma cells on our SCope platform and the ATAC-seq data on a UCSC hub to jointly serve as a resource for the melanoma field. Together, this exhaustive analysis of melanoma cell state diversity indicates that additional states exists between the two extreme melanocytic and mesenchymal-like states. The GRN we identified may serve as a new putative target to prevent the switch to mesenchymal cell state and thereby, acquisition of metastatic and drug resistant potential.

## Introduction

The degree of heterogeneity in cancer is extremely high, and occurs at multiple levels. Solid tumors represent complex mixtures of various cell types, including more than 50 host stromal cell types and cancer cells (Lambrechts et al. 2018). These cancer cells also frequently differ substantially, even within a given tumor, both genetically (e.g. copy number variations) and phenotypically (e.g. transcriptional profile), jointly referred to as intratumoral heterogeneity. In addition, different tumors can differ significantly, even in the same patient, representing an additional layer of intertumoral heterogeneity. Single-cell technologies such as scRNA-seq have the potential to deconvolve this heterogenous complexity, and to identify common transcriptional states and hence common groups of similar cells across tumors. Mostly however, cancer cells from each sample form a distinct cluster per patient, whereas the corresponding normal host cells from various patients cluster together according to their cell type (Tirosh et al. 2016; Puram et al. 2017; Lambrechts et al. 2018). This observation is somewhat counterintuitive because cells with similar gene expression profiles are known to occur in multiple tumors, for instance cells in specific cell cycle stages. In fact, gene regulatory network inference using SCENIC has been shown to normalize away part of these tumor-specific differences, resulting in one pan-tumor cluster of cycling cells (Aibar et al. 2017). Nonetheless, the unsupervised discovery of common transcriptional states remains a challenge.

Melanoma skin cancer is notorious for its pronounced heterogeneity as a result of its high number of irreversible genetic alterations (Alexandrov et al. 2013) and its elevated cellular plasticity (Grzywa, Paskal, and Wlodarski 2017). The latter dynamic process, commonly referred to as phenotype switching (Hoek and Goding 2010), involves reversible transcriptional changes and emerges from the underlying epigenome (Verfaillie et al. 2015), similarly to the epithelial-to-mesenchymal transition (EMT) in other cancers. Initially, two main transcriptional states were found to reoccur across tumors and cohorts of patient-derived cultures (Hoek et al. 2006, 2008; Eichhoff et al. 2011; Zipser et al. 2011; Landsberg et al. 2012; Caramel et al. 2013; Sun et al. 2014; Konieczkowski et al. 2014; Denecker et al. 2014; Müller et al. 2014; Wouters et al. 2014; Verfaillie et al. 2015; Riesenberg et al. 2015). The melanocyte-like state displays high levels of lineage-specific transcription factors, including SOX10 and MITF, and functional pathways associated with the lineage, such as pigmentation. The mesenchymal-like state on the other hand shows expression of SOX9 and activity of AP-1, and has acquired increased migratory and invasive potential and resistance to targeted and immunotherapy. Recently, scattered evidence illustrates the existence of additional, intermediate state(s) (Haass et al. 2014; M. Ennen et al. 2015; Marie Ennen et al. 2017a; Falletta et al. 2017; Tsoi et al. 2018; Rambow et al. 2018; Tuncer et al. 2019).

Here, we perform an exhaustive analysis to investigate the diversity of melanoma cell states and to examine intra- and intertumoral phenotypic heterogeneity. Through the comprehensive profiling of single-cell gene expression (single-cell RNA-seq of 39,263 melanoma cells) and chromatin accessibility (ATAC-seq) in baseline conditions, we compare the gene regulatory networks (GRN) between the melanocytic, mesenchymal, and the previously poorly-described intermediate state. We investigate whether GRNs emerge from a mixture of stable states in the culture, or from a “mixed GRN” that operates in all cells of the culture. Furthermore, to identify key regulators underlying the switch from the melanocytic to mesenchymal state and their importance over time, we align GRNs from different cultures during time-series single-cell RNA-seq profiling. All scRNA-seq data, including SCENIC analyses, are available online as .*loom* files on our single-cell analysis platform SCope (http://scope.aertslab.org/#/Wouters_Human_Melanoma); the ATAC-seq data is available through a UCSC track hub.

## Results

### Despite inter- and intra-tumoral heterogeneity, melanoma cultures group into distinct cell states

To study the diversity of melanoma cell states, including any potential intermediate state(s), we performed scRNA-seq on a cohort of melanoma samples: nine patient-derived 2D cultures (“MM lines”; (Gembarska et al. 2012; Verfaillie et al. 2015)) and one long-term melanoma cell line (A375). (**Supplementary Tables S1,2**). First, we multiplexed all cultures into a single 10x Chromium lane (**Figure 1a**), followed by computational demultiplexing using culture-specific SNPs (**Supplementary Note 1**) (SNPs were derived from previously-published bulk RNA-seq data (Verfaillie et al. 2015)), yielding on average 432 cells per culture (**Supplementary Table S1**). Importantly, this reduces the risk of batch effects while retaining the biological differences between the samples. Note that we also performed 10x on individual MM lines (see further below and **Supplementary Note 1**), which confirmed MM line-specific transcriptomes after demultiplexing. Analogously to scRNA-seq performed on biopsies in any cancer type (Tirosh et al. 2016; Venteicher et al. 2017; Puram et al. 2017; Lambrechts et al. 2018), each cancer sample has a unique transcriptome and forms a distinct cluster after dimensionality reduction (t-SNE; **Figure 1b**). Nevertheless, these melanoma cultures express many common genes, such as markers for cell cycle and for their cells-of-origin, here melanocytes (**Figure 1c-e** and **Supplementary Figure S1a,b**). All MM lines (but not A375) express *TFAP2A* (**Figure 1c**), a known marker of the neural border, the progenitor domain of the neural crest during development (de Crozé, Maczkowiak, and Monsoro-Burq 2011), that is also expressed in melanocytes and melanoma cells (Seberg et al. 2017). In agreement with previous work (Hoek et al. 2006; Verfaillie et al. 2015; Shakhova et al. 2015), the MM lines fall into two main transcriptional states, one showing *SOX10* expression and the other *SOX9* (**Figure 1d**). SOX10, an established lineage transcription factor in melanocytes together with MITF (Southard-Smith, Kos, and Pavan 1998), is expressed in seven of the melanoma cultures (MM001, MM011, MM031, MM057, MM074, MM087 and A375). The remaining three cultures display expression of *SOX9* (MM029, MM047 and MM099; **Figure 1d**). SOX9 is pivotal for the migration of early neural crest cells (Cheung and Briscoe 2003), and for the determination of the chondrogenic cell lineage from the neural crest (Mori-Akiyama et al. 2003), and is believed to represent a marker for melanoma cells that have undergone a state transition, referred to as phenotype switching, towards a de-differentiated, mesenchymal-like and therapy-resistant cell state (Hoek and Goding 2010; Tsoi et al. 2018).

**Figure 1:**
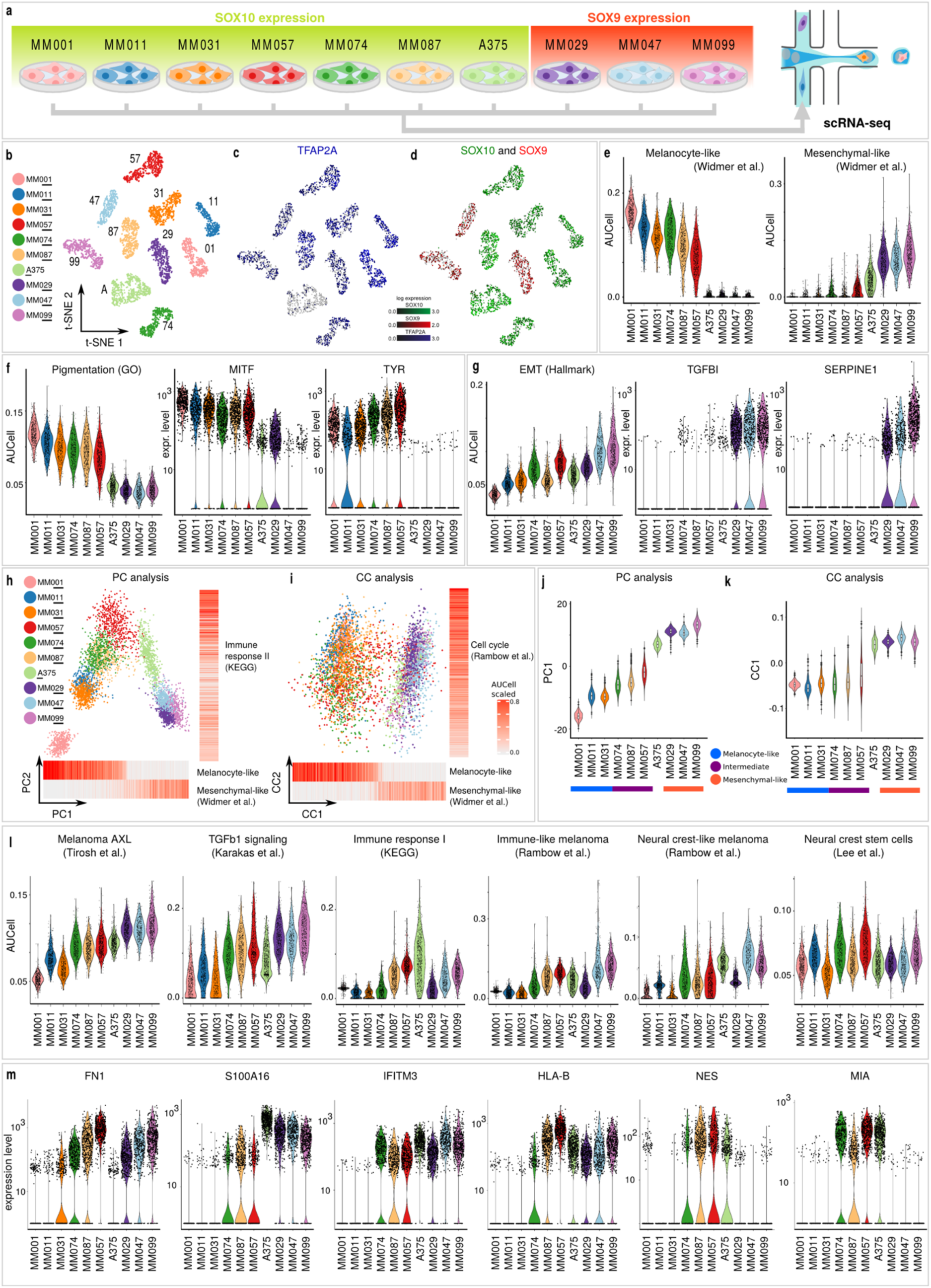
While showing inter- and intra-tumoral heterogeneity, melanoma patient samples cluster into three distinct subtypes. a, Nine patient-derived MM lines and one commercial cell line (A375) were multiplexed into a single 10x Chromium lane, followed by computational demultiplexing using SNPs. **b**, Cells cluster according to their cell line origin in a t-SNE. **c**, All cell lines except A375 express the neural border marker *TFAP2A*. **d**, By the expression of *SOX10* and *SOX9*, the ten cell lines split into melanocyte-like (SOX10-positive) and mesenchymal-like (SOX9-positive) melanomas. **e**, Public melanocyte-like and mesenchymal-like gene signatures (Hoek et al. 2006; Widmer et al. 2012) define two groups among the ten cell lines, with A375, MM029, MM047 and MM099 having a mesenchymal-like transcriptional phenotype. **f**, The melanocyte-like cell lines show high expression of a pigmentation gene signature (GO), and genes involved in melanogenesis, *MITF* and *TYR*. **g**, The mesenchymal-like cell lines have higher expression of a gene signature for epithelial-to-mesenchymal transition (Hallmark) (Liberzon et al. 2015), and MM029, MM047 and MM099 express the epithelial-to-mesenchymal transition genes *TGFBI* and *SERPINE1*, while they are merely expressed in the other cell lines. **h-i**, Both principal component (h) and canonical component (i) analysis order the cells in a gradient from melanocyte-like to mesenchymal-like on the first axis, as illustrated by AUCell activity (see also Supplementary Figure S1c,d). The second axis correlates with immune response (principal component 2) and cell cycle (canonical component 2). **j-k**, The intermediate cell lines show higher mean and variance in PC1 loading (j), and higher variance in CC1 loading (k), compared to the other melanocyte-like cultures (for the boxplots center line represents the median; box limits are upper and lower quartiles). **l**, The melanoma AXL program (Tirosh et al. 2016), and TBFb1 signaling (Karakas et al. 2006), increase in expression from purely melanocyte-like to intermediate to mesenchymal-like cell lines. The intermediate cell lines show higher expression than the melanocyte-like cell lines in immune activation genes (KEGG graft versus host disease, (Kanehisa and Goto 2000), immune- and neural crest-like melanoma genes (Rambow et al. 2018), and neural crest stem cell genes (G. Lee et al. 2007). **m**, The mesenchymal-like genes *FN1* and *S100A16*, along with the immune-related genes *IFITM3* and *HLA-B* are highly expressed in the intermediate and mesenchymal-like cell lines, while the neural crest stem cell markers *NES* and *MIA* are specific to the intermediate cell lines.

When these two groups are contrasted against each other, the differentially expressed genes correspond to previously described signatures of melanocyte-like versus mesenchymal-like transcriptional cell states (**Figure 1e** and **Supplementary Figure S1a**) (Hoek et al. 2006; Widmer et al. 2012; Verfaillie et al. 2015). The SOX10-positive cultures are melanocyte-like, with high expression of pigmentation-related genes, as shown by Gene Ontology analysis (GO), and further illustrated by the expression of MITF, and its target gene TYR, an enzyme essential for melanin production (**Figure 1f**). The SOX9-positive cultures on the other hand show increased expression of genes involved in the epithelial-to-mesenchymal transition, such as TGFBI and SERPINE1 (**Figure 1g**). In addition, the SOX9-positive MM lines show differential expression of genes up-regulated in melanoma cells that have acquired resistance to pharmaceutical BRAF inhibition after prolonged exposure to the drug (Shaffer et al. 2017), and genes specific for melanomas resistant to anti-PD-1 immunotherapy (Hugo et al. 2016) **(Supplementary Figure S1a**), corroborating a phenotype switch. Finally, as expected, each melanoma culture contains subpopulations of cells in specific phases of the cell cycle (**Supplementary Figure S1b**).

To identify additional common sources of variation across the melanoma samples, we performed principal component analysis (PCA) as well as canonical correlation analysis (CCA), using Seurat (Butler et al. 2018) (**Figure 1h-m**). The first principal component corresponds to the distinction between *SOX10*-positive (melanocyte-like) and *SOX9*-positive (mesenchymal-like) states, while high PC2 represents a transcriptional state related to the immune response, and to a lesser extend also to neural crest stem cells, adding an additional layer of inter-individual heterogeneity, mainly within the melanocyte-like cell state (**Figure 1h,j**). Gene set enrichment analyses (GSEA) on PC1 confirms significant enrichment of multiple melanocyte-like (negative PC1) and mesenchymal-like (positive PC1) gene sets (**Supplementary Figure S1c**). CCA removes even more sample-specific differences, with CC1 corresponding to the SOX10/SOX9 cell state axis, and CC2 to cell cycle and translation (**Figure 1i,k, Supplementary Figure S1d**). Interestingly, PCA and CCA also reveal differences in the *degree* of PC/CC loading between MM lines, with a higher variance in MM074, MM087 and MM057 compared to other the melanocyte-like cell lines. Together, these analyses indicate the existence of at least two subtypes of melanocyte-like cell states, with MM074, MM087, and MM057 (from now on referred to as the intermediate MM lines) having increased mesenchymal-like (PC1) *and* immune-response-like (PC2) properties when compared to MM001, MM011 and MM031 (the exclusively melanocyte-like MM lines, from now on referred to as melanocyte-like; **Figure 1h-k**). Indeed, these transcriptional differences are clearly visible when retrospectively zooming in on these specific biological processes. A stepwise increase is detected for the melanoma AXL program (Tirosh et al. 2016) and TGFb1 signaling (Karakas et al. 2006) (**Figure 1l**). The intermediate cultures exhibit increased activity of immune response genes and high resemblance to immune-like melanoma cells in patient-derived xenograft tumors upon combined pharmaceutical BRAF/MEK inhibition (Rambow et al. 2018), and to neural crest stem cells (Rambow et al. 2018; G. Lee et al. 2007) (**Figure 1l**). Examples of specific genes with higher expression across all intermediate compared to melanocyte-like samples include *FN1* and *S100A16* (mesenchymal-like), *IFITM3* and *HLA-B* (immune-related) and *NES* and *MIA* (neural crest stem cell marker) (**Figure 1m**). Interestingly, the intermediate samples show a higher *variance* in PC1/CC1 loading, indicating higher intra-sample heterogeneity for MM074, MM087, and MM057 (**Figure 1h-k**), superimposed on their increase in “mean” PC1/CC1 loading. Importantly, this increased heterogeneity is observed in both PC and CC analyses, but is more distinct when using CCA. Thus, CCA is suitable to remove patient-specific bias and discover underlying common expression programs as well as heterogeneity of transcriptional states.

We also quantified heterogeneity at the gene level, by calculating the Gini coefficient for each gene within each sample, using GiniClust (Jiang et al. 2016; Tsoucas and Yuan 2018). A Gini coefficient of 0 means that all cells show the same level of expression, while 1 means that a single cell expresses all the mRNA molecules and the other cells express none. GSEA on these Gini gene rankings demonstrates heterogeneous expression of mesenchymal-like genes in a subset of melanoma cultures, i.e. MM031, MM074, MM087 and MM057 (**Supplementary Figure S2a**). To verify these observations, we performed Drop-seq on a biological replicate of MM057 cells (Macosko et al. 2015) (**Supplementary Table S1]**). Interestingly, MM057 cells in Drop-seq occupy the same area in the CCA plot as 10x MM057 cells (**Supplementary Figure S2b**). In addition, there is a high correlation between Gini coefficients of both techniques (**Supplementary Figure S2c**), indicating that the heterogeneity is stable over time and independent of the experimental technique used for scRNA-seq.

In conclusion, by using scRNA-seq on a panel of nine MM lines, three levels of heterogeneity can be identified, namely: (1) inter-individual differences between samples; (2) differences between subtypes or cell states; (3) intra-individual differences, with different degrees of variation.

### Melanoma transcriptional cell state predicts single-cell migratory capacity

Because the three observed cell states are determined by differences in mesenchymal-like and neural crest-related genes, we wanted to test whether these transcriptional differences become manifest in a relevant phenotype, such as migration. Therefore, we investigated the migratory capacity of the melanoma samples at single-cell level. We seeded cells on a collagen-coated microfluidic chip (Mathieu et al. 2016), containing a large seeding channel connected to 70 microchannels (6 x 10 µm) on both sides in which the cells can migrate (**Figure 2a**). Because the microchannels are smaller than the cell’s diameter, only individual cells fit, allowing to study their migratory behavior in 1D at single-cell level. We performed live-cell imaging of the patient-derived melanoma cells over a period of 24 hours, taking an image every 4 minutes using a lens-free imaging device (Mathieu et al. 2016). In lens-free imaging, objectives are replaced with lasers, to produce digital holograms, and images are digitally reconstructed. This has several advantages over conventional live-cell imaging platforms such as space reduction, a larger field of view and the possibility for *a posteriori* focusing (Isikman et al. 2011).

**Figure 2:**
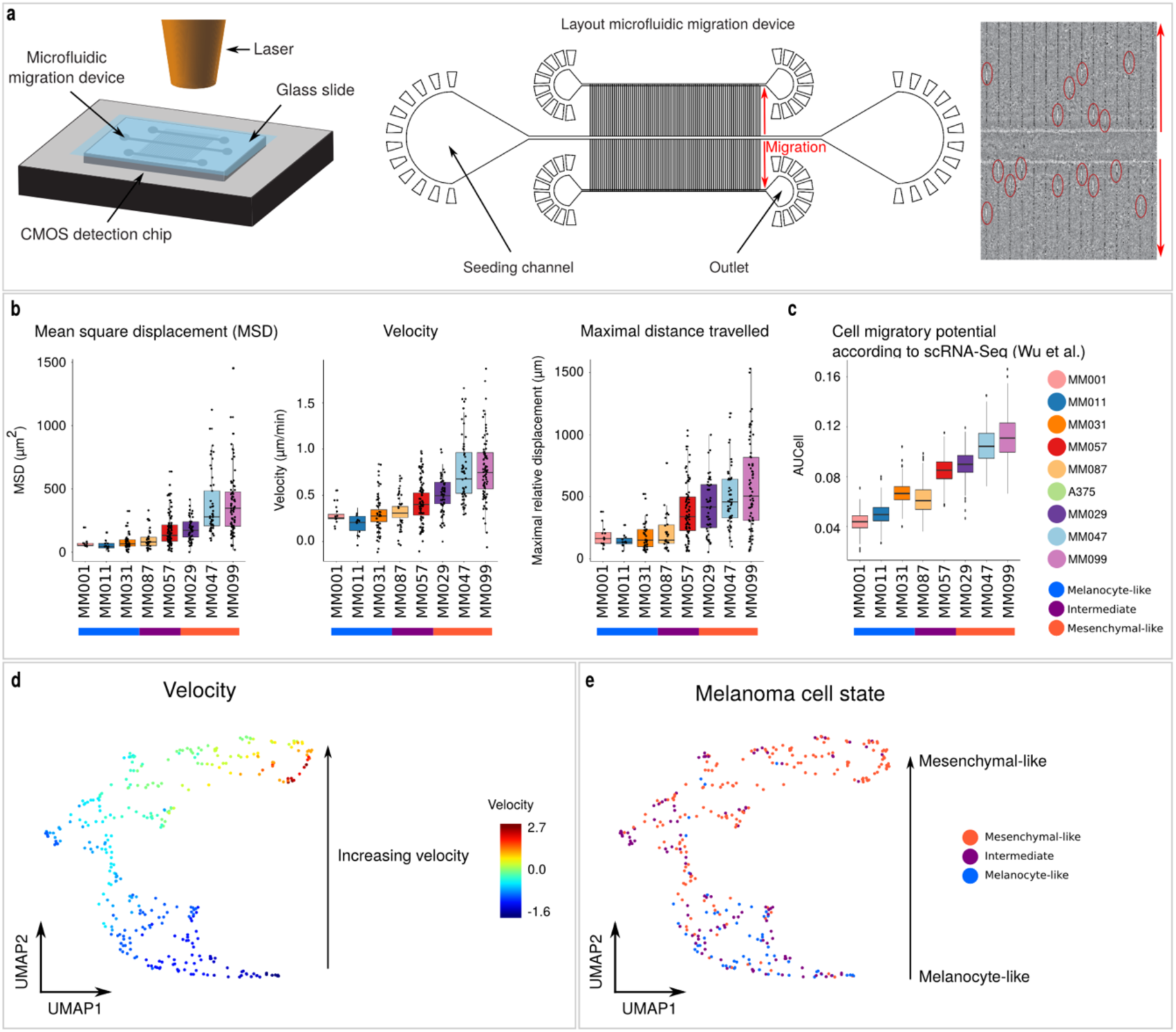
The melanoma transcriptional cell state predicts single-cell migratory capacity. **a**, Single-cell migration assay set-up using lens-free imaging. Left: A microfluidic device containing single-cell migration channels is loaded with cells, covered with a glass slide and placed on top of a complementary metal-oxide-semiconductor sensor (CMOS) that detects diffraction of the emitted laser though the migration device to track the movement of single cells. Middle: Layout of the microfluidic migration device. Cells are loaded in the seeding channel and will migrate through the small migration channels. Right: Reconstructed image of cells (encircled) moving in the migration channels in the direction of the red arrows. **b**, The mean square displacement (left), velocity (middle) and maximal distance travelled (right) of single cells show a gradual increase from melanocyte-like to mesenchymal-like cell lines (center line, median; box limits, upper and lower quartiles). **c**, The expression of cell migratory potential genes (Wu et al. 2008) correlates with the migratory behavior lines (center line, median; box limits, upper and lower quartiles). **d-e**, UMAP with single cells based on their migratory capabilities colored by velocity (d) and by cell state (e).

Tracking individual cells over time for 24 hours enabled calculating their mean squared displacement, a common metric to calculate the diffusion of particles or cells over time (see methods). The mean squared displacement revealed a wide variety of migratory capacity within and across melanoma cultures (**Figure 2b**). The three *SOX9*-positive MM lines (MM029, MM047 and MM099) cover the longest distance within 24 hours, in line with the transcriptional resemblance of these cultures to migratory mesenchymal cells. Interestingly, two of the three heterogeneous cultures with an intermediate cell state, namely MM057 and MM087, also show a high degree of cell migration (**Figure 2b**). The third intermediate sample, MM074, was phenotypically different in cell shape when attached to collagen in the device as compared to the normal incubation surface (plastic), and was removed from these analyses. A similar picture is obtained by calculating the average velocity and maximal distance reached (**Figure 2b**). Strikingly, a gene signature predictive for the rate of cellular migration in bladder cancer (Wu et al. 2008) is relevant and applicable to melanoma as well, i.e., it follows the same global trend as the empirically measured migration (comparing **Figure 2b** and **2c**; see also **Supplementary Figure S3**). To get a global overview of the cells across all melanoma cultures, we combined various physical features of migratory behavior (mean squared displacement, velocity and maximal distance) and performed dimensionality reduction, using UMAP (Becht et al. 2018), on the cell-feature matrix. Interestingly, cells do not cluster per culture but follow a continuous gradient that strongly correlates with velocity (**Figure 2d**). Mesenchymal-like melanoma cell lines are enriched among the fastest-migrating cells, whereas melanocyte-like and intermediate melanocyte-like cell lines are enriched among the cells that migrate slowest and with intermediate velocity, respectively (**Figure 2e**).

In conclusion, using a single-cell migration assay, we confirmed that the mesenchymal-like melanoma cultures are highly migratory cell lines. Whereas the heterogeneous and intermediate cell state patient-derived melanoma cultures display intermediate migratory potential.

### Single-cell network inference reveals candidate regulators of the intermediate cell state

To further examine the predicted melanoma cell states, we applied SCENIC network inference to the single-cell expression matrix (Aibar et al. 2017). SCENIC predicts transcription factors (TFs) governing each melanoma cell state, alongside candidate transcription factor target genes. A transcription factor with its candidate targets is called a regulon. SCENIC yields a regulon-cell matrix with regulon activities across all single cells, and provides therefore an alternative dimensionality reduction. A UMAP visualization based on the regulon-cell matrix reveals three candidate cell states in an unsupervised manner, recapitulating our findings above (**Figure 3a**). One cluster represents the mesenchymal-like cell state (MM029, MM047 and MM099); while the remaining SOX10-positive samples can be divided between a fully melanocyte-like (MM001, MM011, MM031) and an intermediate cell state (MM074, MM087 and MM057). The intermediate state shares several regulons with the melanocyte-like cell state such as SOX10, MITF, IRF4, SOX4 and USF2 (**Figure 3b**, for a full list of detected regulons see **Supplementary File 1**). The exclusively melanocyte-like cell state also displays activity of the HES6 regulon. Note that while most of these melanocyte-like regulons have a discrete activity pattern, MITF shows a gradual decrease in activity (**Figure 3b** and **Supplementary Figure S4a**). The intermediate state also shares regulons with the mesenchymal-like cell state, such as AP-1 members (JUN, FOSL2, FOSB, FOSL1) and important immune modulators IRF/STAT (**Figure 3b,c**). The latter observation is in line with our previous observation that these cultures have elevated expression of genes characterizing immune-like melanoma cells (**Figure 1l**). Finally, some regulons are specific to the intermediate state, including EGR3, RXRG and NFATC2 (Figure 3b,c). These TFs have previously been linked to a more aggressive/dedifferentiated phenotype in cancer and/or in melanoma specifically (Baron et al. 2015; Aibar et al. 2017; Rambow et al. 2018), which confirms that these cultures are in-between the melanocyte-like and the more deleterious mesenchymal-like state (Hoek et al. 2006; Verfaillie et al. 2015). Of note, cell-line-specific TFs were also detected (such as SOX11 in A375), and each melanoma culture contains cells with increased activity of cell cycle regulons such as FOXM1 (**Figure 3b**).

**Figure 3:**
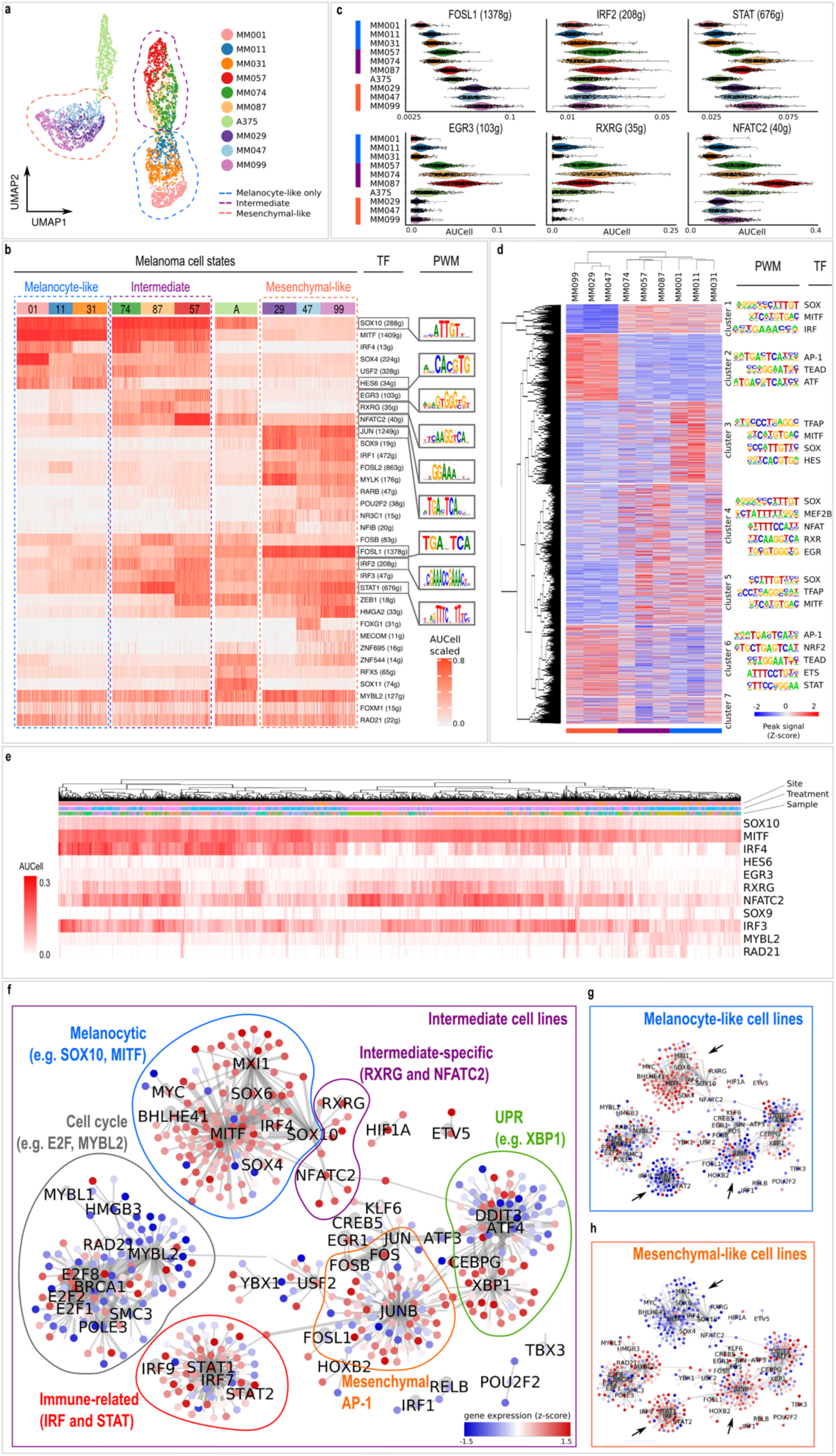
Single-cell network inference reveals candidate regulators of the intermediate cell state. **a, A** SCENIC-based UMAP separates unsupervised the intermediate cultures from the other melanocyte-like cultures. **b**, The heatmap shows a selection of the regulons identified by SCENIC (rows), and their activities in each cell (columns). For each regulon, the transcription factor (TF) and the number of predicted target genes is indicated. For selected TFs the DNA-binding motif is shown, predicted by the RcisTarget step of SCENIC (for a complete list of the SCENIC regulons see Supplementary File 1). **c**, Violin plots showing the activity of SCENIC regulons specific for the intermediate cultures (bottom) and shared between the intermediate and the mesenchymal-like cultures, as measured by AUCell for all melanoma cultures. **d**, Chromatin accessibility, measured by Omni-ATAC-seq, in the MM lines, establishes three cell line clusters, corresponding to the previously described cell states, and seven clusters of genomic regions. For each cluster of regions, the enriched DNA-binding motifs and associated TFs are indicated. **e**, Heatmap showing the activity of selected regulon (rows) in each cell (columns) as identified by SCENIC in a publicly available dataset of 2,018 single cells isolated from 32 patient melanomas (Tirosh et al. 2016; Jerby-Arnon et al. 2018). As reported previously (Aibar et al. 2017), the majority of cells is melanocyte-like (activity of the SOX10 and MITF regulon) with few cells being fully melanocyte-like (activity of HES6 regulon) and most cells intermediate (activity of EGR3, NFATC2 and RXRG). **f-h**, Gene regulatory networks as identified by SCENIC across melanoma cultures, displaying gene expression in the intermediate (left), melanocyte-like (top right) and mesenchymal-like (bottom right) MM lines (relative to rest) as node color. The edge width corresponds to the number of SCENIC runs in which the TF-target interaction is predicted. The network consists of several substructures (transcriptional programs) indicated by the colored lines. Arrows indicate transcriptional programs that differ between melanoma cell states.

Next, to validate our findings, we profiled the chromatin landscape of these nine MM lines, using Omni-ATAC-seq (Corces et al. 2017). As expected, the melanocyte- and mesenchymal-like cultures display preferential accessibility of the previously-identified state-specific H3K27Ac regions (Verfaillie et al. 2015) (**Supplementary Figure S4b**). In line with the transcriptome data, we observe one mesenchymal-like and two melanocyte-like groups of melanoma cultures, and identify clusters of accessible regions for each of them (**Figure 3d**; see Methods). Genomic regions accessible in both groups of melanocyte-like cultures (cluster 1 and 5) are indeed enriched for the SOX10, MITF and IRF4 binding motifs, whereas those in the mesenchymal-like cultures (cluster 2) are characterized by AP-1 and TEAD motifs. This is confirmed by the observation that SOX10-bound regions from two public SOX10 ChIP-seq datasets (Laurette et al. 2015; Eskiocak et al. 2017), are accessible in the melanocyte-like cultures, but not in the mesenchymal-like ones (**Supplementary Figure S4c**). In addition, NFATC2, RXRG and EGR3 motifs are enriched in the open regions of the intermediate samples (cluster 4), and the HES6 motif in the other melanocyte-like cultures (MM001, MM011 and MM031; cluster 3). Of note, even though both MITF and HES6 bind to E-box motifs, the two observed motifs differ substantially (-CATGTGAC- for MITF and -CACGTG- for HES6), and correspond between scRNA- and Omni-ATAC-seq data. Similar to our observations in the scRNA-Seq data, open regions shared between the intermediate and the mesenchymal-like cultures are enriched for the AP-1 and IRF/STAT motifs (cluster 6; **Figure 3d**). This is further corroborated by public AP-1 ChIP-seq data (Gertz et al. 2013; Joseph et al. 2010), in which AP-1-bound regions display highest accessibility in mesenchymal-like and intermediate cultures (**Supplementary Figure S4c**). Interestingly, the AP-1 peaks that are more accessible in the intermediate cultures are observed in the proximity of marker genes of these intermediate cultures, such as FN1, IRF2, NFATC2 and SOX9 (**Supplementary Figure S4d**). Next, we investigated the existence of the intermediate cell state in human clinical melanomas. First, we verified a publicly available data set of 2,018 malignant single cells isolated from 32 patient melanomas, predominantly metastases (1,896 of 2,018 cells) (Tirosh et al. 2016; Jerby-Arnon et al. 2018). As shown previously (Aibar et al. 2017), most cells in this cohort display activity of the SOX10 regulon (**Figure 3e**). Few of them seem to reside in a fully melanocyte-like melanoma cell state, indicated by an overall high HES6 regulon activity. The majority of the cells from this public data set are in the intermediate cell state, showing activity of EGR3, NFATC2 and/or RXRG. This is in line with the elevated migratory capacity of the intermediate cell state and the fact that those cells originate mainly from metastases. Indeed, metastatic melanoma cells display higher activity of those intermediate regulons (Jerby-Arnon et al. 2018) (**Supplementary Figure S4e**). Importantly, melanomas with high activity of one of the regulons also have elevated activity of the others (**Supplementary Figure S4f**), indicating that these tumors indeed represent an alternative, intermediate cell state (this correlation is not observed when comparing to the HES6 regulon; **Supplementary Figure S4f**). Note that the correlation between these three regulons is not because of high overlap of their target genes (not a single gene is common to all three regulons). Second, we also confirmed the intermediate cell state in *The Cancer Genome Atlas* (TCGA) bulk RNA-seq cohort of 375 patient melanomas (Cancer Genome Atlas Network 2015). Melanoma samples exist in this cohort that show high activity of the intermediate regulons (NFATC2, RXRG, EGR3) and their corresponding target genes (**Supplementary Figure S4g**). Again, high correlation is observed between the activity of all three regulons, whereas there is an inverse correlation with the activity of the HES6 regulon. And finally, the intermediate cell state is also observed in Omni-ATAC-seq data of TCGA tumors (Corces et al. 2018) (**Supplementary Figure S5**). Interestingly, the regions associated with the intermediate and mesenchymal-like cultures seem to be more generally accessible across different cancer types as observed by others (Rambow et al. 2015), whereas the regions linked to melanocyte-like cultures are highly specific for melanoma skin cancer, and the intermediate cell state-linked enhancers are also accessible in gliomas.

In conclusion, gene regulatory network analysis of the scRNA-seq data shows that melanoma cultures MM074, MM087 and MM057 do not only represent heterogeneous cell populations but also an alternative transcriptional melanoma cell state sharing characteristics of the melanocyte- and mesenchymal-like cell states (**Figure 3f-h**). Importantly, this intermediate cell state is also detected in corresponding ATAC-seq data, and in melanoma biopsies.

### SOX10 perturbation leads to recurrent state transitions

Next, we wanted to investigate whether dynamic cell state *transitions* are also heterogeneous, and to what extent they are influenced by the baseline cell state. To this end, we knocked-down (KD) *SOX10* in the six melanocyte-like cultures (MM001, MM011, MM031, MM074, MM087 and MM057). *In vivo*, a decrease in *SOX10* expression occurs upon acquiring resistance to targeted therapy (BRAF and/or MEK inhibition; (Sun et al. 2014; Shaffer et al. 2017)) and immunotherapy (Landsberg et al. 2012). Therefore, and because of the central role for SOX10 in the melanocyte-like cell state (Verfaillie et al. 2015), we hypothesised that SOX10-KD would reprogram melanocyte-like cells towards the SOX9-positive, mesenchymal-like cell state. Firstly, SOX10-KD resulted in varying degrees of cell death, with the lowest cell death in the intermediate samples (MM074, MM087 and MM057) and the highest cell death in the melanocyte-like cultures (MM001, MM011 and MM031; data not shown). Bulk RNA-seq 72 hours after SOX10-KD shows an extensive down-regulation of the melanocyte-like lineage markers in all cultures, as expected, and in addition, up-regulation of the mesenchymal-like gene signature to a varying degree (**Figure 4a**). To investigate this process in higher resolution, we performed scRNA-seq after knockdown of SOX10, and knockdown with a non-targeting control siRNA, and for technical reasons focussed on the melanoma cultures with the lowest degree of cell death (MM057, MM074 and MM087; see **Figure 4b** for experimental setup). In order to unravel the dynamic aspects of the state transition, we included multiple time points (24, 48 and 72h), and in addition included a mixture of cells sampled every 2 to 4 hours (2, 4, 6, 8, 12, 16, 20, 24, 28 and 32 hours) for two of the three cultures (MM057 and MM087). Visualization of all cells simultaneously and coloring according to the experimental time point clearly indicates the trajectory that melanoma cells follow after SOX10-KD, regardless of the method for dimensionality reduction (t-SNE or Diffusion Maps; **Figure 4c**), enabling us to calculate a pseudotime for every single cell (see Methods). Interestingly, using cellAlign (Alpert et al. 2018) we confirmed that a highly similar trajectory is inferred by specialized trajectory inference methods, such as SCORPIUS (Cannoodt et al. 2016) and Monocle 2 (Qiu et al. 2017) (**Supplementary Figure S6**). Down-regulation of *SOX10* and up-regulation of *SOX9* accompanies this trajectory (**Figure 4d**), reaffirming the antagonistic roles of the two transcription factors. As expected, established target genes of SOX10, including *MITF, DCT* and *TYR*, decrease in expression over time and in fact lag behind *SOX10* expression (**Figure 4e**). Although the three melanoma cultures display slightly distinct dynamics (in MM087 cells, the expression of SOX10 targets drops at around 24 hours, between 24 and 48 hours in MM057, and after 48 hours in MM074 cells), the expression changes between the three cultures are highly concordant. This could be demonstrated by applying dynamic time warping, using cellAlign (Alpert et al. 2018) (**Figure 4f)**. Thus, the state transition after SOX10-KD is a recurrent phenomenon across cultures, and may therefore reflect a specific genomic regulatory program.

**Figure 4:**
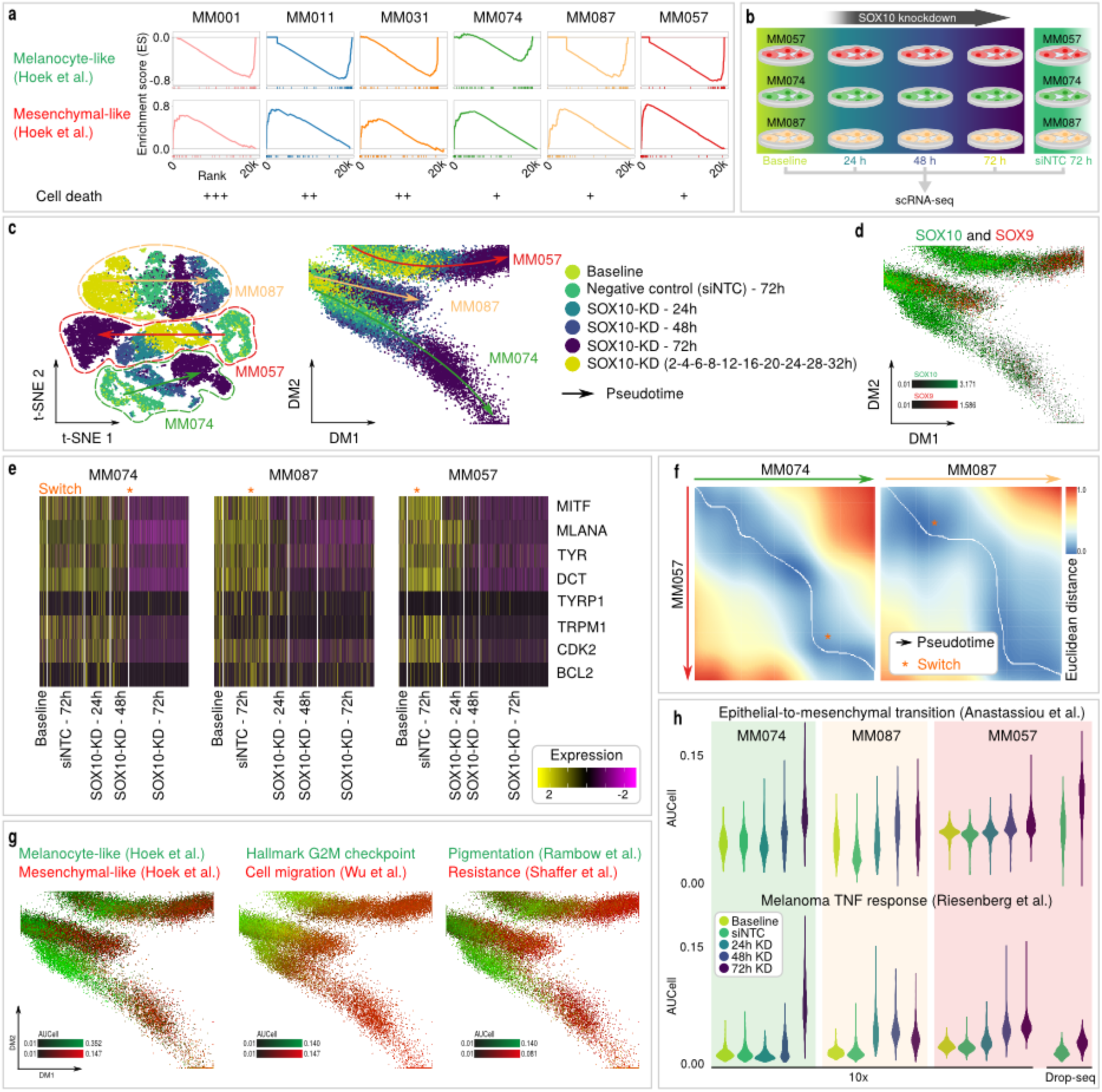
Transcription factor perturbation reveals recurrent state transitions. **a**, Gene Set Enrichment Analysis plots of the melanocyte-like (top) and mesenchymal-like (bottom) signatures (Hoek et al. 2006; Widmer et al. 2012) for each MM line 72 hours after knockdown (KD) of SOX10 (relative to 72 hours after knockdown with a non-targeting control siRNA). Semi-quantitative assessment of cell death is indicated at the bottom (‘+++’: high cell death; ‘++’: intermediate; ‘+’: low cell death). **b**, Experimental setup. **c**, Seurat t-SNE (left) and Diffusion Map (MP; right) with cells colored according to the experimental time point, indicated the trajectory that melanoma cells follow after SOX10-KD (also see colored arrows). **d**, While *SOX10* (green) is down-regulated along the trajectory from control to 72h SOX10-KD, *SOX9* is upregulated (red). **e**, Expression of established SOX10 target genes (rows) in each cell (columns) grouped by experimental time point. The three cultures display slightly distinct dynamics (see orange asterisk). **f**, Comparative alignment of transition trajectories by applying dynamic time warping, shows the optimal alignment (white line) through a dissimilarity matrix. Despite the varying dynamics (orange asterisk indicate the switch from melanocytic to mesenchymal transcriptional program), the expression changes between the three cultures are highly concordant. **g**, Seurat MP with cells colored by expression of melanocyte-like (green) and mesenchymal-like (red) genes (Hoek et al. 2006; Widmer et al. 2012) measured by AUCell for all melanoma cultures (left), of hallmark G2M checkpoint (green) and cell migration (red) (Wu et al. 2008) genes (middle), and of pigmentation (green) (Rambow et al. 2018) and therapy resistance (red) (Shaffer et al. 2017) genes (right). **h**, Violin plots showing the expression of the epithelial-to-mesenchymal transition (top) (Anastassiou et al. 2011), and of the melanoma TNF response (bottom) (Riesenberg et al. 2015) measured by AUCell for all melanoma cultures. For MM057, 10x Chromium and Drop-seq (on biological replicate cells) scRNA-seq data is shown.

The magnitude of the transcriptional switch is extensive, with a collapse of the cell cycle and the entire melanocytic transcriptional program (see methods; **Figure 4g,h** and **Supplementary Figure S7b-d**). An average of 6,337 genes shows decreasing expression within 72 hours after SOX10-KD (6,495, 6,396 and 6,113 in MM057, MM074 and MM087, respectively; **Supplementary Figure S7a**), whereas 1,369 genes exhibit up-regulated transcription (1,209, 1,308 and 1,591, respectively). Analogous to the common set of down-regulated genes after SOX10-KD, the up-regulated genes are largely shared between melanoma cultures, too (see Methods; **Supplementary Figure S7b**). The up-regulated processes involve cellular migration, the epithelial-to-mesenchymal transition, cancer metastasis, immune cell activation, angiogenesis, and melanoma-specific gene sets such as the signatures for the SOX9-positive cultures (Hoek et al. 2006; Verfaillie et al. 2015), the melanoma TNF response (Riesenberg et al. 2015), the AXL program signature (Tirosh et al. 2016) and the previously-mentioned signature for acquired resistance to BRAF inhibition (Shaffer et al. 2017) (**Figure 4g,h** and **Supplementary Figure S7b-d**). To examine the validity of these results independent of the experimental technique used for scRNA-seq, we also performed Drop-seq (Macosko et al. 2015) on MM057 melanoma cells before and 72 hours after SOX10-KD. These data validate the consistency of the observed transcriptional changes after SOX10-KD, i.e., the disruption of cell cycle and the melanocytic program and the resulting induction of gene sets including migration, the epithelial-to-mesenchymal transition, SOX9-positive cultures and acquired resistance to BRAF inhibition (**Supplementary Figure S7d)**.

In conclusion, using scRNA-seq, we demonstrate that SOX10-KD in melanocyte-like cultures reprograms cells to the mesenchymal-like cell state during phenotype switching. The switch is highly reproducible across cultures, displays varying dynamics depending on the beginning state, and involves a complete collapse of cell cycle and of the lineage transcriptional program.

### Single-cell network inference reveals the sequence of recurrent dynamic gene regulatory changes during phenotype switching

Next, to predict dynamic gene regulatory changes after SOX10-KD, we applied SCENIC network inference along the trajectories’ pseudotime (Aibar et al. 2017). In total we detected 477 regulons, uniformly divided over the three melanoma cultures with 269, 254 and 281 regulons identified for MM074, MM087 and MM057, respectively (for all regulons, see **Supplementary File 1**). Approximately a quarter of them are shared across all three melanoma cultures (23.9% or 114 of 447). These results confirm the recurrence of the state transition and allow us to focus on the dynamic gene regulatory changes during phenotype switching (for all common regulons, see **Supplementary Figures S8-10**). The initial event after knockdown of *SOX10* is the pausing of cell cycle, as demonstrated by the inactivation of members of the E2F and MYB transcription factor families, and of DNA polymerase (**Figure 5a** and **Supplementary Figures S11** for the gene regulatory networks over time). Almost simultaneously, there is an initial increase in the unfolded protein response (UPR; e.g. XBP1) and in AP-1 activity (e.g. JUNB). Afterwards, the melanocyte core program is shut down, as evidenced by a loss of *SOX10* and *MITF* transcriptional activity. Additional transcription factors that are paused at this time include CEBPZ, MYC and ETS-related factor ETV5. Simultaneously, there is an induction of ATF/CREB (e.g. CREB3), AP-1 (e.g. FOSB and JUN) and ETS-related factor ELK1, and a further increase of the previous UPR and AP-1 factors. Finally, immune-related transcription factors IRF/STAT (e.g. IRF7/9 and STAT1) become activated. Of note, each melanoma culture also exhibits the involvement of culture-specific transcription factors, such as MEF2A, NFE2L1 and RELA for MM074, NR2F2 and RARG for MM087 and NR3C1 and RARB for MM057.

**Figure 5:**
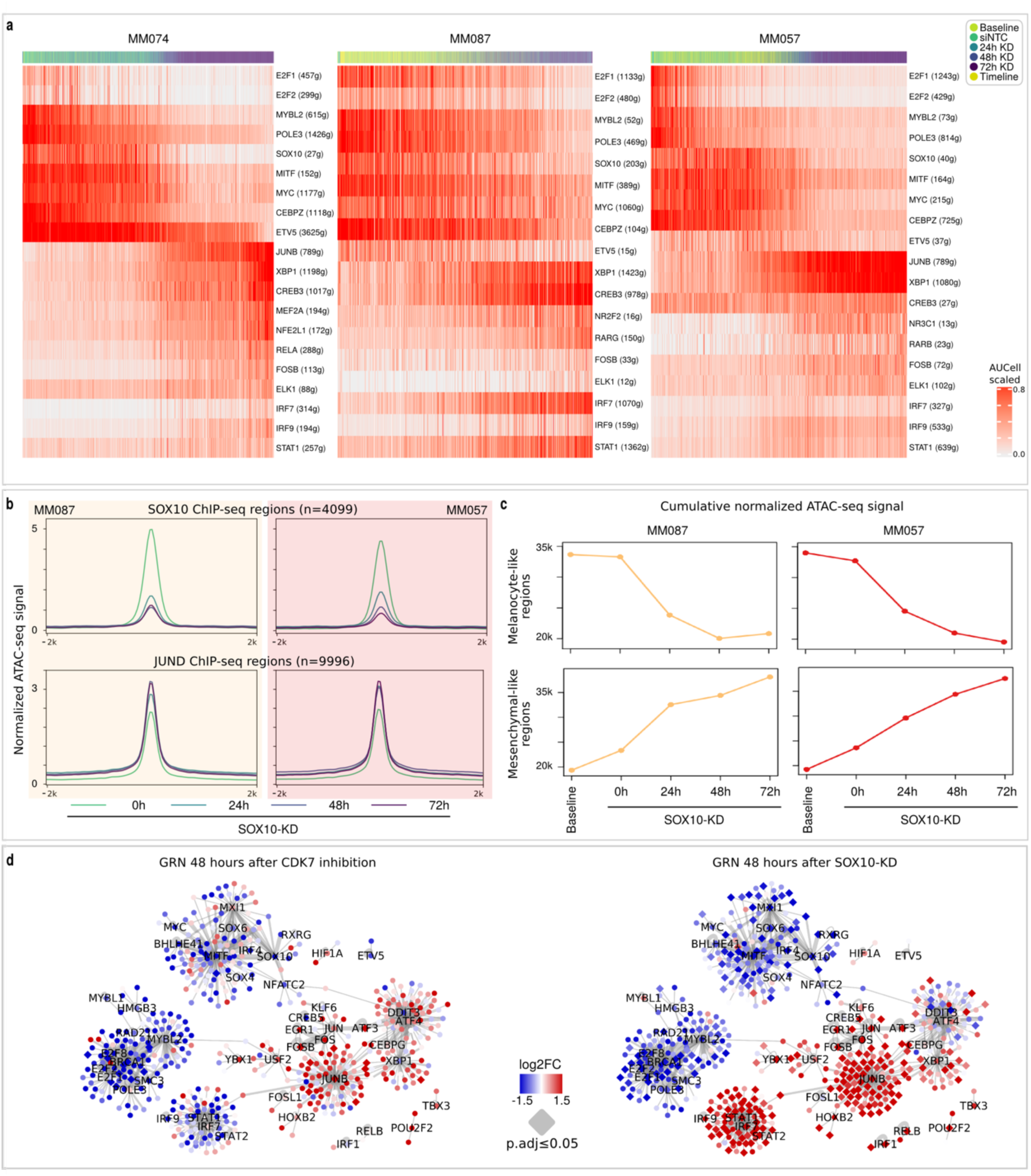
Single-cell network inference the reveals sequence of recurrent dynamic gene regulatory changes during phenotype switching. **a**, Heatmap showing the regulon (rows) activities in each cell (columns) as identified by SCENIC. Cells are ordered according to their diffusion maps’ pseudotime. (see also Figure 4c,f). For each regulon, the transcription factor (TF) and the number of predicted target genes is indicated. **b**, Normalized ATAC-seq signal in SOX10-bound regions (top) and JUND-bound regions (bottom), as previously identified by ChIP (Laurette et al. 2015; Gertz et al. 2013). **c**, Cumulative normalized ATAC-seq signal in melanocyte-like regions (top) and mesenchymal-like regions (bottom), as shown in Figure 3d (melanocyte-like, cluster 1; mesenchymal-like, cluster 2). **d**, Gene regulatory networks (GRNs) as identified by SCENIC across melanoma cultures (see also Figure 3f-h), displaying gene expression 48 hours after CDK7 inhibition by THZ2 (left) and after SOX10-KD (right). genes that change significantly (p.adj<=0.05) are shown in diamond shape. The edge width corresponds to the number of SCENIC runs in which the TF-target interaction is predicted.

To validate these observations we analyzed Omni-ATAC-seq data on the same experimental conditions for two of the melanoma cultures (MM087 and MM057; Bravo González-Blas et al. 2019). Analogous to our observations at the transcriptome level, a global collapse of the melanocytic chromatin landscape is observed, as exemplified by the closure of SOX10-bound regions (**Figure 5b**) (Laurette et al. 2015) and of melanocyte-like regions (**Figure 5c**; cluster 1 in **Figure 3d**). These epigenome changes occur already 24 hours after SOX10-KD and hence precede the transcriptional switch (**Figure 5b** versus **Figure 4e-f**). On the other hand there is a gradual increase in accessibility of mesenchymal-like regions (**Figure 5c**; cluster 2 in **Figure 3d**) and of AP-1-bound regions (**Figure 5b**) (Gertz et al. 2013). Because of the observed collapse of both the melanocytic transcriptome and epigenome, we hypothesized that SOX10-KD could be mimicked by a general inhibition of transcription. Therefore, we treated the three melanoma cultures (MM074, MM087 and MM057) with THZ2, a selective inhibitor of CDK7 (Wang et al. 2015), and sampled cells for bulk RNA-Seq 6 and 48 hours later. To verify efficient treatment, we investigated the expression of genes previously found down-regulated in melanoma cells as early as 6 hours after CDK7 inhibition (using THZ1, a parent compound) (Eliades et al. 2018). Indeed, the expression of these genes is down-regulated 6 and 48 hours after the start of the treatment (**Supplementary Figure S12**), confirming effective CDK7 inhibition. Next, we compared the transcriptional changes of the melanoma cells’ GRN after THZ2 treatment to those after SOX10-KD (both at 48 hours). The loss in activity of the melanocytic and cell cycle transcriptional programs is indeed recapitulated by a general inhibition of transcription (**Figure 5d**). Also the enhanced activity of the UPR and AP-1 programs are mirrored by CDK7 inhibition. Unlike after SOX10-KD, no increase in the immune-related IRF/STAT transcriptional program is observed.

In conclusion, using dynamic GRN inference, we unravel the sequential arrangement of transcriptional programs during phenotype switching that ultimately lead to the mesenchymal-cell state. In addition, we demonstrate that CDK7 inhibition partially recapitulates SOX10-KD-induced phenotype switching.

## Discussion

Here, we present a comprehensive study of intra- and intertumoral heterogeneity of regulatory melanoma cell states, both in steady-state conditions and during dynamic cell state transitions. We have made all scRNA-seq data, including SCENIC analyses, available online on our in-house developed single-cell analysis and visualization tool SCope (http://scope.aertslab.org/#/Wouters_Human_Melanoma) and all ATAC-seq data on a UCSC track hub. This will markedly facilitate the application of these data in other research studies, including in the area of cancer cell states, gene expression variation, and melanoma cell biology.

Using scRNA-seq data of the steady-state melanoma cultures, we detected established melanoma cell states, specifically the melanocyte-like (or proliferative) and the mesenchymal-like (or invasive) cell state, and confirmed the corresponding active transcriptional programs. Mesenchymal-like cells have lost most of their lineage identity, and have acquired programs responsible for migration and resistance to immune and targeted therapy (Wu et al. 2008; Hugo et al. 2016; Shaffer et al. 2017). Importantly, we also functionally validated the mesenchymal-like cell’s increased migratory potential in terms of distance and speed in a single-cell migration assay.

The higher resolution of scRNA-seq compared to bulk RNA-seq enabled us to uncover and study intratumoral heterogeneity within cultures classified as either of these two cell states. Its unsupervised discovery however is considerably complicated by the predominating intertumoral heterogeneity. Integration and comparison of single-cell datasets across samples is a focus of attention in the bioinformatics field, as multiple methods have been suggested to remove batch effects or warp one data set onto another (Haghverdi et al. 2018; Welch et al. 2019; Stuart et al. 2019; Luecken and Theis 2019). Here, we found that side-by-side comparison of different dimensionality reduction techniques (PCA, CCA, and regulatory network inference by SCENIC) provide an interesting angle to interpret heterogeneity.

Besides the expected heterogeneity in cell cycle phases, we discover mesenchymal-like cells within the intermediate melanocyte-like cultures. These cells vary in their *degree* of mesenchymal phenotype, with some of them being equally potent as cells in mesenchymal-like cultures. The mesenchymal-like cultures on the other hand represent a more homogenous population of cells with only cell cycle generating heterogeneity. Interestingly, we show that the heterogeneity is stable over time and across replicates of the same culture, and can be detected regardless of the scRNA-seq technology applied. This strongly suggests it to be a regulated rather than a stochastic process (Foreman and Wollman 2019). Using single-molecule RNA FISH for a selected panel of marker genes, Shaffer and colleagues observed a similar rare, semi-coordinated transcription in other melanoma cultures (Shaffer et al. 2017). Using scRNA-seq, we here demonstrate that these rare cells can be identified in an unsupervised manner at the whole-transcriptome level, enabling the discovery of novel marker genes and processes.

The existence of the melanocyte-like and mesenchymal-like cell state is well-established (reviewed in Rambow et al., *in press*) (Arozarena and Wellbrock 2019). Already at the time of the discovery of these cell states, a group of cell lines was observed that share characteristics of both states, and that were therefore left out of the classification, the so-called cohort B cell lines (Hoek et al. 2006; Widmer et al. 2012). Consequently, melanoma cells were observed that also combine functional properties of both cell populations, i.e., invading melanoma cells that retain their proliferation capacity (Haass et al. 2014; Falletta et al. 2017). Recently, comprehensive studies discovered alternative cell states, notably cells that have acquired a transcriptional profile reminiscent of neural crest stem cells (Tsoi et al. 2018; Rambow et al. 2018). In addition, this research also detected cell lines that combine properties of the two main cell states, now SOX10 and AXL (Tsoi et al. 2018), or MITF and AXL expression (Tuncer et al. 2019). While it could *a priori* have been conceivable that ‘semi-invasive’ melanoma cell lines (based on bulk RNA-seq) would consist of a mixture of cells in either the melanocyte-like or the mesenchymal-like state, our results now establish that these intermediate transcriptomes are largely due to a stable “mixed gene regulatory network” that produces this intermediate transcriptome in all cells, and only partly due to heterogeneity between the cells or random flipping between the states. Recently, Foreman and Wollman analogously demonstrated that expression variability in mammary epithelial cells (MCF10A) results from cell state differences rather than transcriptional bursting (Foreman and Wollman 2019). Notably, these observations were made using MERFISH instead of scRNA-seq. Using SCENIC, we found indeed gene regulatory networks (GRNs) that are specific for this intermediate state. Besides those specific networks, the intermediate state shares GRNs with the melanocyte and mesenchymal states, recapitulating previous observations (Hoek et al. 2006; Widmer et al. 2012; Haass et al. 2014; Falletta et al. 2017; Tsoi et al. 2018; Tuncer et al. 2019). Activity of most melanocytic lineage transcription factors, including SOX10 and MITF, and corresponding functional activities, such as pigmentation, are retained, albeit frequently at a lower degree. On the other hand, the cells in this state have acquired a set of mesenchymal-like properties, and share a subset of master regulators of that cell state, most notably IRF/STAT and some of the AP-1 family members. The activity of these transcription factors is reflected by the cells’ transcriptional and functional profile, i.e. an activated immune system phenotype and an increased migratory potential, respectively.

The specific master regulators found in the intermediate network include EGR3, NFATC2 and RXRG. Interestingly, these transcription factors have been implicated in mechanisms that could explain the intermediate cell state’s characteristics. EGR3, as a member of the early growth response genes, is needed for the induction of important cellular programs such as neurogenesis (Kim et al. 2012; Quach et al. 2013; Meyers et al. 2018), inflammation and the immune response (S. Li et al. 2012; Baron et al. 2015), VEGF-mediated angiogenesis (Liu Dan et al. 2003; Kang et al. 2014) and the maintenance of stemness (Hamra et al. 2004). In line with these observations, gene set enrichment analysis (GSEA) of the SCENIC-predicted EGR3 target genes shows enrichment of gene signatures characteristic for vasculature development and stem cells. Interestingly, SCENIC identifies EGR3 as the master regulator of the intermediate cell state, predicting that it controls the other two transcription factors NFATC2 and RXRG (while neither NFATC2 nor RXRG are predicted to regulate EGR3). NFATC2 is an established modulator of the immune system (Peng et al. 2001; Nguyen et al. 2010; Walters et al. 2013), and has been implicated in the dedifferentiation of melanoma cells (Perotti et al. 2016; Aibar et al. 2017) and the aggressive behaviour of cancer cells in general (Baumgart et al. 2012; Griesmann et al. 2013). GSEA of the NFATC2 targets indeed demonstrates similar functions, including the wounding response, the epithelial-to-mesenchymal transition, and stemness. In general, NFATC2 is believed to induce transcription, but multiple reports indicate a function as transcriptional repressor as well (Ranger et al. 2000; Baksh et al. 2002; Carvalho et al. 2007; Baumgart et al. 2012). And finally, RXRG is a member of the ligand-responsive transcription factor family of retinoic acid/X receptors and is able to activate a variety of cellular processes, depending on its partner nuclear receptor in the DNA-binding heterodimer complex (Altucci et al. 2007; de la Fuente et al. 2015). It promotes the dedifferentiation and invasion of tumor cells (Liu et al. 2011; Papi et al. 2010) and has been shown to be involved in minimal residual disease of melanoma (Rambow et al. 2018). GSEA of the RXRG targets indeed indicates corroborating functions such as the maintenance of stemness, neural crest stem cells, oligodendrocyte markers and myogenesis. Comparative analysis of our RXRG target genes and those identified by Rambow et al. (Rambow et al. 2018) highlights distinct sets (only 4 shared genes of 109 in total), which may be due to the presence of alternative heterodimer partners. The intermediate cell state displays high expression of MITF, in contrast to the RXRG-positive cells responsible for minimal residual disease (Rambow et al. 2018). We previously identified AXL as a marker for mesenchymal-like cultures, and did not detect it as a marker for our intermediate cell state. This, in combination with the observation that the intermediate state displays high activity of immune response transcriptional programs, strongly suggests that it represents yet another cell state, distinct from the AXL/SOX10-positive (Tsoi et al. 2018) and MITF/AXL-positive cells (Tuncer et al. 2019). Of note, as is clear from the SCENIC UMAP (**Figure 3a**), both the melanocyte-like and intermediate cell state represent a spectrum of GRNs. In fact, each melanoma culture has its own unique transcriptome of which some do show AXL expression (the mesenchymal-like cultures MM029, MM047 and MM099; the intermediate culture MM087; and the A375 cell line).

Importantly, we confirm the existence of the intermediate cell state in melanoma biopsies. In a single-cell RNA-seq cohort of 32 patient samples and 2,018 melanoma cells (Jerby-Arnon et al. 2018), we detected most melanoma cells to be in the intermediate state. Because most of these cells are known to be metastatic cells, this observation is in line with the cell state’s aggressive and dedifferentiated phenotype and increased migratory capacity. A previous study, describing single-cell qPCR data of 472 cells of primary melanomas (n=5), confirms the existence of malignant cells expressing both MITF-high and MITF-low signature genes (Marie Ennen et al. 2017b), but was not able to identify the master regulators of these cells because of the limited set of analyzed genes. Interestingly, we could also reveal the presence of the intermediate state in bulk genomics data of large clinical cohorts (TCGA), both from RNA- and ATAC-seq data. Altogether, this makes these specific transcription factors attractive candidates for targeted melanoma therapy, or cancer in general. NFATC2 and RXRG have been suggested and assessed before as potential therapeutic targets in melanoma (Perotti et al. 2016; Rambow et al. 2018). Because of its role as master regulator, EGR3 is likely to be the best candidate for therapeutic targeting. In addition, in a different context, EGR3 inhibition has been shown to block AP-1 activity upon NGFR activation (Levkovitz and Baraban 2002).

*SOX10* and its target genes are responsible for the lineage specification and differentiation of many neural-crest derived cell types, including melanocytes and the corresponding cancer melanoma (Simões-Costa and Bronner 2015; Verfaillie et al. 2015). Yet, melanoma cells adopt a range of cell state phenotypes showing varying degrees of differentiation, with the most extreme cell state having lost most of its lineage identity and conversely showing a mesenchymal-like or de-differentiated phenotype (Hoek et al. 2006; Verfaillie et al. 2015; Rambow et al. 2018). Here, we demonstrate that knockdown of SOX10 is sufficient to switch melanocyte- to mesenchymal-like melanoma cells, and that the resulting cells acquire the same transcriptional programs for migration, invasion and resistance to therapy as mesenchymal-like cells (Wu et al. 2008; Verfaillie et al. 2015; Hugo et al. 2016; Shaffer et al. 2017). This observation strongly suggests that *SOX10* directly or indirectly suppresses the mesenchymal-like cell state and the associated gene regulatory networks. Although *SOX10* has been suggested as a transcriptional repressor in breast cancer (Dravis et al. 2018), we could not detect any evidence for such a role in melanoma (f.i., there is no enrichment of SOX binding sites near genes with up-regulated expression after SOX10-KD). Alternatively, given the large-scale effects of SOX10-KD on gene expression, the phenotype switch likely results from the loss of *SOX10*-controlled repressors, including transcriptional and post-translational repressors. In fact, the emergence of the mesenchymal-like cell state is presumably the combined result of multiple down-regulated repressors, and might involve *SOX10* target genes DUSP6, MXI1 and NFATC2 (Schreiber-Agus et al. 1995; T. C. Lee and Ziff 1999; Carvalho et al. 2007; Wong et al. 2012; Qu et al. 2012). We also demonstrate that the inhibition of CDK7-dependent transcription mimics the extensive effects of SOX10-KD on gene expression, leading to the inhibition of cell cycle and the melanocytic program, but also the increasing activity of the AP-1 program and the unfolded protein response. This suggests that the up-regulation of the AP-1 program can at least in part be due to a stress response, resulting from the collapsing transcriptome. On the other hand, genes up-regulated after SOX10-KD include SOX9 and other markers from the mesenchymal-like melanoma state, arguing against a pure stress response signature. Notably, this observation warrants further research into the usage of such molecules in a clinical setting.

In conclusion, we used single-cell transcriptomics combined with gene regulatory network and trajectory inference to map the gene regulatory landscape of recurrent melanoma cell states. We find that transcriptional and phenotypic heterogeneity can be largely attributed to differences in gene regulatory networks, supplemented with some degree of stochasticity; and that phenotype switching from the melanocytic to the mesenchymal state is controlled by a regulatory program that is highly reproducible across distinct patient cultures.

## Material and methods

### Cell culture

The melanoma cultures (MM001, MM011, MM029, MM031, MM047, MM057, MM074, MM087 and MM099) are derived from patient biopsies by the Laboratory of Oncology and Experimental Surgery (Prof. Dr. Ghanem Ghanem) at the Institute Jules Bordet, Brussels (Gembarska et al. 2012; Verfaillie et al. 2015). They can be obtained from Dr. Ghanem Ghanem with a Material Transfer Agreement. Cells were cultured in Ham’s F10 nutrient mix (ThermoFisher Scientific) supplemented with 10% fetal bovine serum (Invitrogen) and 100 µg ml-1 penicillin/streptomycin (ThermoFisher Scientific). The human melanoma cell line A375 was obtained from the ATCC and were maintained in Dulbecco’s Modified Eagle’s Medium with high glucose and glutamax (ThermoFisher Scientific), supplemented with 10% fetal bovine serum (Invitrogen) and penicillin-streptomycin (ThermoFisher Scientific). Cell cultures were kept at 37°C, with 5% CO2 and were regularly tested for mycoplasma contamination, and were found negative. Knockdown of SOX10 was performed using a SMARTpool of four siRNAs (SMARTpool: ON-TARGETplus SOX10 siRNA, number L017192-00, Dharmacon) at a final concentration of 20nM in Opti-MEM medium (Thermo Fisher Scientific). To control for transfection artefacts (‘siNTC’), we used a pool of non-targeting siRNAs (SMARTpool: ON-TARGETplus Non-targeting Pool, number D001810-10-05, Dharmacon). SOX10-KD and siNTC treatments of MM001, MM011, MM031, MM047, MM057 and MM087 were sampled after 72h for bulk RNA-seq. SOX10-KD treatments of MM047, MM057 and MM087 were sampled after 24, 48 and 72h for single-cell RNA-seq and Omni-ATAC-seq (MM057 & MM087 only). Additionally, MM057 and MM087 were sampled after 2h, 4h, 6h, 8h, 12h, 16h, 20h, 24h, 28h and 32h SOX10-KD, and treatments were pooled per cell line (‘timeline’ sample) for 10x scRNA-seq. Inhibition of CDK7-dependent transcription was achieved by treatment with THZ2 for 6 and 48h (HY-12280, MedChemExpress) (Wang et al. 2015). For each treated melanoma culture, IC50 concentrations were calculated for 48 hours of treatment. As a control, DMSO treatment was used.

### Single-cell migration

The single-cell migration devices were fabricated in a clean room using a standard soft lithography process, as described by (Tong et al. 2012). In short, master silicon wafers were patterned with the design of the microfluidic chip as shown schematically in **Figure 2a**. As this was a two-layered design (seeding channel height 25 µm and migration channels 10 µm height), SU-8 2025 and SU-8 2010 (MicroChem) were used sequentially. The PDMS devices were produced by mixing the PDMS prepolymer and crosslinker in a weight ratio of 10:1, followed by de-gassing the PDMS in vacuum for 45 minutes before pouring it onto the silicon wafers. The PDMS was cured for 2 hours at 80°C, after which the cured devices were carefully peeled from the wafer and cut to size. Inlet and outlets were created using a 3 mm-diameter biopsy puncher (Electron Microscopy Sciences). PDMS chips were bonded onto a cleaned glass glide using plasma. Just before use, collagen type I (Thermofisher Scientific) was incubated in the channels for 1 hour at room temperature, followed by rinsing twice with DPBS and twice with medium. In the meantime, confluent cells were washed with DPBS, detached, spun down at 1,000 RPM for 5 minutes and resuspended in medium. The cells were seeded in the coated and rinsed microfluidic chip by adding 10 µl of the cell solution in one of the inlet wells and, after a few seconds, adding the same volume of cells into the other inlet well. To let the cells attach to the seeding channel, the chip was incubated for 20 minutes at 37°C. Once the cells were attached, the inlet wells were rinsed with medium and each inlet and outlet well was filled with 10 µl of medium. The chip was covered with a glass slide to avoid evaporation and was placed on the CMOS sensor of the lens free imaging (LFI) device (Mathieu et al. 2016) in the incubator (see **Figure 2a** for set-up). Samples were imaged every 4 minutes over a period of 24 hours. The recorded holograms were digitally reconstructed by choosing to focal depth at which the cells were optimally visible inside migration channels. The resulting images were stacked into a video, which was used to manually track the single cells over time using ImageJ. Using the time (*t*) and time-dependent coordinate data (*y*(*t*)), the maximal distance travelled (*y*_*max*_), the mean squared displacement (*MSD*) and the velocity (*v*) per single cells, were calculated as following:

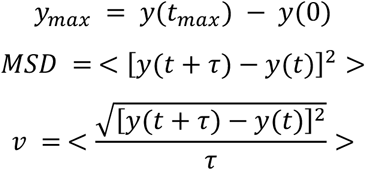

Where <…> stand for time averaging, *t*_*max*_ for the maximal time (24 hours), and *τ* for the time lag, i.e. *τ* = *n* * *dt* with *dt* being the time between two consecutive frames (4 minutes) and *n* the considered time step (*n* = 1 for the velocity and *n* = 5 for the *MSD*).

### Bulk RNA-seq

#### RNA extraction

Total RNA was extracted from attached cells at the indicated time points using the innuPREP RNA mini kit (Analytik Jena), according to the manufacturer’s instructions. Quality assessment was performed using the Bioanalyzer 1,000 DNA chip (Agilent) after which libraries were established.

#### Library preparation and sequencing

For the SOX10-KD samples, total RNA was enriched for mRNA using the Dynabeads mRNA purification kit (Invitrogen). To make cDNA, 1 µl of oligo(dT) primers (500ng/µl; Ambion) and 1 µl of 10 mM dNTP (Promega) was added to 10 µl of polyA-selected mRNA; incubated at 65°C for 5 minutes and placed on ice. First-strand cDNA synthesis was performed by adding 4 µl of first strand buffer (Invitrogen), 2 µl of 100 mM DTT (Invitrogen) and 1 µl of Superscript II (Invitrogen) and incubating the mix at 42°C for 50 minutes, then 70°C for 15 minutes. The second strand of cDNA was filled in by adding 35 µl of water, 15 µl of 5x second strand buffer (Invitrogen), 1.5 µl of 10 mM dNTP, 0.5 µl of 10 U/µl E Coli DNA ligase (Bioke), 2 µl of 10 U/µl E Coli DNA polymerase I (Bioke), 1 µl of 2 U/µl E Coli RNaseH (Invitrogen) and then incubating at 16°C for 2 hours. The cDNA was purified on a MinElute column (Qiagen) and eluted in 15 µl EB buffer. To incorporate sequencing adapters, we combined the purified cDNA with 4 µl of Nextera TD buffer (Illumina) and 1 µl of Nextera Tn5 enzyme (Illumina) on ice and incubated at 55°C for 5 minutes. The tagmented cDNA was purified again on a MinElute column and eluted in 20 µl EB buffer. To PCR amplify the fragments, we added 25 µl of NEBnext PCR master mix (Bioke), 5 µl of Nextera primer mix and incubated at 72°C for 5 minutes, then at 98°C for 30 sec, followed by 15 cycles of 98°C for 10 sec, 63°C for 30 sec and 72°C for 3 minutes. We purified the PCR amplicons with 55 µl AMPure beads (Analis). Final libraries were pooled and sequenced on a NextSeq500 (Illumina).

For the THZ2 samples, RNA-seq libraries were prepared according to the Illumina Truseq stranded mRNA sample preparation guide. Final libraries were pooled and sequenced on a HiSeq4000 (Illumina).

#### Sequencing data processing

RNA-seq reads were cleaned with fastq-mcf from the ea-utils package (v1.04.807) (Aronesty [2015] 2011, 2013) and mapped to the genome (hg19) using STAR (v2.5.1b) (Dobin et al. 2013). Read counts per gene were obtained from the aligned reads using htseq-count (v0.6.1p1) (Anders, Pyl, and Huber 2015). The Bioconductor/R package DESeq2 (v1.18.1) (Love, Huber, and Anders 2014) was used for normalization and differential gene expression analysis.

#### Data analysis

Log2FoldChange values were used for ranking the genes, and downstream Gene Set Enrichment Analysis (GSEA) (Subramanian et al. 2005).

### scRNA-seq

#### Cell preparation

Cells were washed, detached by trypsinization, spun down at 1,000 RPM for 5 minutes to remove the medium, resuspended in PBS-BSA, filtered through a 40 μm cell strainer, counted and processed according to the downstream protocols.

#### General sequencing data processing

The single cell libraries were sequenced on HiSeq4000 and NextSeq500 instruments (Illumina), most libraries were sequenced twice. Sequencing parameters can be found in **Supplementary Table S2**). RNA-Seq data quality was assessed with FastQC (v0.11.5) (Babraham Bioinformatics n.d.) and MultiQC (v1.0.dev0) (Ewels et al. 2016).

#### 10x Genomics: Library preparation

Single-cell libraries were generated using the GemCode Single-Cell Instrument and Single Cell 3′ Library & Gel Bead Kit v2 and Chip Kit (10x Genomics) according to the manufacturer’s protocol. Briefly, melanoma cells were suspended in 0.04% BSA–PBS. About 8,700 cells were added to each channel with a targeted cell recovery estimate of 5,000 cells. After generation of nanoliter-scale Gel bead-in-EMulsions (GEMs), GEMs were reverse transcribed in a C1000 Touch Thermal Cycler (Bio Rad). After reverse transcription, single-cell droplets were broken and the single-strand cDNA was isolated and cleaned with Cleanup Mix containing DynaBeads (Thermo Fisher Scientific). cDNA was then amplified with a C1000 Touch Thermal Cycler. Subsequently, the amplified cDNA was fragmented, end-repaired, A-tailed and index adaptor ligated, with SPRIselect Reagent Kit (Beckman Coulter) with cleanup in between steps. Post-ligation product was amplified with a C1000 Touch Thermal Cycler. The sequencing-ready library was cleaned up with SPRIselect beads (see also **Supplementary Table S2**).

#### 10x Genomics: Sequencing data processing

The 10x Chromium single cell gene expression data were generated with seven sequencing runs (see **Supplementary Table S2**). Each sequencing run was processed with 10x Genomics companion software CellRanger for alignment, barcode assignment and UMI counting (using the reference set hg19-1.2.0 provided by 10x Genomics). The number of cells in each sample was estimated by the CellRanger cell detection algorithm. Filtered count matrices were converted to sparse matrices in R using Seurat (v2.1.0) (Butler et al. 2018), and cells expressing less than 1,000 genes and cells with more than 20% mitochondrial reads were removed. The Seurat objects of the sequencing runs were then merged and meta data were populated using treatment and cell line ID (with demuxlet resolved barcode-to-sample ID assignment in case of mixed sequencing runs; **Supplementary Note 1**). This resulted in a dataset of 32,738 genes by 43,178 cells. For quality metrics see **Supplementary Table S1**.

#### Drop-seq: Library preparation

Single-cell libraries were generated as previously described (Macosko et al. 2015), according the Drop-seq laboratory protocol (version 3.1, 12/28/2015) with minor modifications. Briefly, melanoma cells were suspended in 0.01% BSA–PBS at a final concentration of 100 cells/µl. Single cells were co-encapsulated with barcoded beads resuspended in lysis buffer (ChemGenes cat. no. Macosko-2011-10) using an Aquapel-coated PDMS microfluidic device with an identical flow rates for both solutions. After processing approximately 1 ml of cell suspension, droplets were broken with perfluorooctanol in 30 ml of 6× SSC. Next, the beads were washed and resuspended in a reverse transcriptase mix, followed by a treatment with exonuclease I to remove unextended primers. The beads were then washed again, counted, aliquoted into PCR tubes, and PCR amplified. The resulting PCR reactions were pooled and purified (first using the minElute PCR purification kit of Qiagen, followed by an extra purification step using AMPureXP beads of Beckman Coulter), and the amplified cDNA quantified on a BioAnalyzer High Sensitivity Chip (Agilent). The cDNA (50 ng) was fragmented and amplified for sequencing with the Nextera DNA sample prep kit (Illumina) using custom primers that enabled the cDNA sequence from Read1 (**Supplementary Table S3**). The libraries were purified, quantified, and then sequenced using spike-in custom primers (see **Supplementary Table S2-3**).

#### Drop-seq: Sequencing data processing

Reads were cleaned for adapters and sequencing primers with fastq-mcf from the ea-utils package (v1.04.807) (Aronesty [2015] 2011, 2013). Drop-seq data were processed using the Drop-seq tools pipeline v1.12 implemented by Macosko et al. 2015 (Macosko et al. 2015; Nemesh [2018] 2019), Picard tools (v1.140) (Broad Institute 2019) and the STAR aligner (v2.5.1b) (Dobin et al. 2013). We selected cell barcodes with at least 1,000 genes expressed, less than 50,000 UMIs and mitochondrial expression percentage below 10. Next, count matrices of the three Drop-seq runs were combined to a matrix of 13,597 genes by 2,278 cells and processed with Seurat (v2.1.0) (Butler et al. 2018) similarly to the 10x data. For quality metrics see **Supplementary Table S1**.

#### Single-cell RNA-seq subsets

We defined the following subsets of the single-cell RNA-seq data: “10 Baselines” (10 cell lines without treatment); “MM057_SOX10-KD” (cell line MM057 baseline sample, no-template control, SOX10-KD [24h, 48h and 72h], SOX10-KD timeline) “MM087_SOX10-KD” (cell line MM087 baseline sample, no-template control, SOX10-KD [24h, 48h and 72h], SOX10-KD timeline) “MM074_SOX10-KD” (cell line MM074 baseline sample, no-template control, SOX10-KD [24h, 48h and 72h]) “MM057_SOX10-KD DropSeq” (Drop-seq on cell line MM057 baseline sample, no-template control, SOX10-KD 72h) “Three_MM_lines_SOX10-KD” (cell lines MM057, MM074 and MM087 baseline sample, no-template control, SOX10-KD [24h, 48h and 72h], and SOX10-KD timeline of MM057 and MM087).

We retrieved those subsets from the full 10x and Drop-seq matrices and analyzed them as follows: raw read counts were CPM-normalized using edgeR (Robinson, McCarthy, and Smyth 2010) and genes that are expressed in less than 1% of the cells were filtered out. Variable genes were identified from the normalized and scaled data, and principal component analysis (PCA) was performed on the expression matrix of the variable genes. t-SNE was performed using the first 10 principal components by default. In some cases, the number of principal components for calculating the t-SNE was selected from the elbow plots generated by Seurat. Diffusion maps were calculated by diffusionMap (v1.1-0.1; Richards 2018) through Seurat’s RunDiffusion function with the option max.dim = 3. The normalized Gini coefficients of the CPM-normalized scRNA-seq data were calculated using GiniClust (Jiang et al. 2016; Tsoucas and Yuan 2018). For the canonical correlation analysis (CCA), we partitioned the “10 Baselines” Seurat object into ten separate culture-specific Seurat objects, and calculated for each of them the top 1,000 variable genes. As final gene list for CCA, we used the union of these genes that are expressed in all ten cultures (3,952 genes). CCA was then performed using the RunMultiCCA function within Seurat (v2.1.0; Butler et al. 2018).

#### Trajectory inference

Three trajectory inference methods was used on the SOX10-KD scRNA-seq data: SCORPIUS (v1.0; Cannoodt et al. 2016), Monocle 2 (v2.6.5; Qiu et al. 2017) and diffusionMap (v1.1-0.1; implemented in Seurat v2.1.0; Richards 2018). For SCORPIUS, we used the normalized expression matrix of variable genes to perform dimensionality reduction with multi-dimensional scaling (MDS) and inferred the trajectory by finding the shortest path between cluster centers obtained by k-means clustering. For Monocle 2, we started with raw UMI counts of genes that are expressed in at least 1% of all the cells. After estimating size factors and dispersion, we performed PCA followed by t-SNE for dimensionality reduction. Clusters were identified in 2D t-SNE space, and top 1,000 genes that are differentially expressed between clusters were used for trajectory inference. For diffusionMap, we used the first three principal components calculated over variable genes as described above. For each culture, we extracted the coordinates of its first two diffusion components and fitted a lowess principal curve, using the princurve R package (Cannoodt [2018] 2019). The arc-length of the fitted curve (lambda) was extracted and used as pseudotime after being scaled to [0,1] range: (x-min(x))/(max(x)-min(x)).

For the comparison of trajectories, we used cellAlign (Alpert et al. 2018). The expression data of each melanoma culture was first interpolated (*interWeight*), with a window size of 0.1 (*winSz*) and 200 desired interpolated points (*numPts*), along its corresponding trajectory inferred by diffusionMap. The resulting matrix was further scaled using the *scaleInterpolate* function. Finally, the interpolated scaled values were used as input to *align* (default parameters: *sig.calc = F, num.perm = 200*) the pseudo times of the cultures in a pairwise fashion (MM057 & MM074, MM057 & MM087). For comparing the trajectories inferred by different methods, we followed a similar approach.

For plotting gene expression along pseudotime, we used the interpolated scaled data across the 200 interpolated points. The genes in the heatmap were selected and ranked using the following procedure. For each gene, we fitted a logit model (y∼phi1/(1+exp(-(phi2+phi3*x)) to the interpolated scaled data from cellAlign. The nonlinear least-squares estimates of the parameters were determined using the nls R package. Next, we removed the genes where no fit was possible. For the remaining genes, we calculated the pseudo-time point where the scaled interpolated expression is halved. This metric was used to rank the genes in the heatmap. We further kept only the genes that were used as input for running SCENIC.

#### Gene Set Activity

The activity of regulons and gene sets in single cells was calculated with AUCell (v0.99.5) (Aibar et al. 2017). AUCell uses the Area Under the Curve (AUC) to calculate whether a given set of genes is enriched within the expressed genes of a cell. First, the algorithm creates a ranking of genes for each cell from highest to lowest expression. Those rankings are then used to build recovery curves with the genes in an input gene set and to calculate the AUC for the top 5% of ranked genes (default value).

To visualize AUCell scores with ComplexHeatmap (v1.17.1) (Gu, Eils, and Schlesner 2016), we scaled the scores for each gene set from 0 to 1. AUCell values visualized as violin and scatter plots were not scaled.

#### Gene Regulatory Network inference with SCENIC

We inferred gene regulatory networks independently on the different subsets of the single-cell RNA-seq data using SCENIC (v0.1.7) (Aibar et al. 2017). Therefore, we log2-transformed the CPM-normalized counts of each subset with a prior of 1 and used those normalized counts to run the coexpression algorithm GENIE3 (Huynh-Thu et al. 2010) implemented in arboreto (v0.1.3) (Moerman et al. 2019), and to subsequently infer gene regulatory networks with SCENIC (using default settings). SCENIC found 324, 281, 254, 269, 175 and 293 regulons in the subsets 10 Baselines, MM057_SOX10-KD, MM087_SOX10-KD, MM074_SOX10-KD, MM057_SOX10-KD Dropseq and Three_MM_lines_SOX10-KD, respectively (**Supplementary File 1**). In each subset, we used the regulon activity represented by AUCell values to cluster the cells with UMAP, using Seurat’s function RunUMAP (with default settings, except for min.dist = 0.2, dims = 1:5, seed.use = 123).

We generated a combined gene regulatory network by merging all SCENIC regulons from the first four subsets. The resulting GRN contains 169,055 connections of 384 TFs to 12,297 target genes. We filtered this network for TF-target connections with a GENIE3 coexpression weight of at least 0.005 that recur in at least two subsets, and obtained a GRN with 910 connections of 146 TFs to 636 target genes. The networks (depicted in **Figure 3f-h**) contain 804 connections of 46 TFs to 533 targets, omitting 100 TFs that do not have recurrent target gene predictions other than themselves. Among those 100 TFs are for example the AP-1 member JUND, that has large regulons in three subsets, but no coexpression weight stronger than 0.003. Or the nuclear receptor ESRRA with regulons with strong coexpression weights in all four subsets, but no recurrent connections across the subsets.

GRNs were visualized using Cytoscape (v3.4.0) (Shannon et al. 2003). To plot gene expression of single-cell data on those GRNs, we first normalized to library size and log2-transformed the aggregated single-cell UMI counts per treatment (**Supplementary Figures S8-10**) or per cell state (**Figure 3f-h**) and z-score normalized each gene using the median expression. Gene expression values of bulk sequencing experiments are displayed as log2-fold change of treated samples compared to control samples (**Figure 5d**).

#### SCope

The single-cell RNA-seq data and the SCENIC results generated in this publication can be explored interactively in SCope (http://scope.aertslab.org/#/Wouters_Human_Melanoma) (Davie et al. 2018). To create the .loom files uploaded to SCope, we used SCopeLoomR (v0.5.0) (aertslab [2018] 2019). Cells are annotated with the metadata ‘Cell_Line’ and ‘Experiment’, and quality metrics can be assessed by visualizing the number of UMIs per cell (‘nUMI’), the number of genes (‘nGene’), and the percentage of mitochondrial reads (‘percent.mito’). We clustered the cells based on expression values (t-SNE, PCA, Diffusion Map, Expression UMAP) and AUCell values of the regulons that were discovered by SCENIC in the respective subset (AUCell UMAP). Those embeddings were added to each .loom file.

#### Ternary plots

For each cell line, we ranked the cells along the diffusionMap pseudotime axis depicted in figure 4c (see Methods ‘Trajectory inference’). Next, we calculated the gene importance for each gene in the logCPM UMI count matrix along this pseudotime, using the function gene_importances of the R package SCORPIUS (v1.0) (Cannoodt et al. 2016) with 100 permutations, sorted the genes according to their importance and calculated the rank ratio. The direction of regulation along the pseudotime axis was determined by contrasting for each cell line the top and bottom 5% of cells with the function FindMarkers of Seurat (v2.1), using the likelihood ratio test (option ‘bimod’) and the options only.pos = F, return.thresh = 1 and logfc.threshold = 0.

The ternary plot in Supplementary Figure 7b displays the rank ratio of pseudotime importance for each cell line, but only for those genes that are present in the importance rankings of all three cell lines and that have a p-value of <=0.05 (as calculated by the gene_importance function of SCORPIUS). Of those 881 genes, 461 are down-regulated and 268 up-regulated in all three cell lines, 152 have a cell-line specific direction of regulation. The colour in the ternary plot corresponds to the mean logFC in those cell lines in which the importance of this gene is significant (p<=0.05). Note that SOX9 is displayed, although it does not have a significant gene importance in any of the cell lines.

To visualize the enrichment of the collection of public gene sets in the three cell lines along the pseudotime axis, we first ranked the genes for each cell line by the logarithm of the SCORPIUS p-value, signed by the direction of regulation given by the logFC. We ran gene set enrichment analysis on those rankings using a collection of public gene sets and the function bulk.gsea of the R package liger (v0.1) (Fan and Kharchenko 2019). The results were filtered for significant gene sets (q.val <= 0.05) and the enrichment scores were scaled from 0 to 1 for visualization in Supplementary Figure S7c. The colour corresponds to the mean enrichment score (NES) of significant enrichments only. The ternary plots were created using the R package ggtern (v3.0.0) (Hamilton and Ferry 2018).

### Omni-ATAC-seq

#### Cell preparation

Cells were washed, detached by trypsinization, spun down at 1,000 RPM for 5 minutes to remove the medium, resuspended in medium, counted and processed according to downstream protocol.

#### Transposition, library preparation & sequencing

Omni-ATAC-seq was performed as described previously (Bravo González-Blas et al. 2019). Briefly, 50,000 cells were pelleted at 500 RCF at 4°C for 5 minutes, medium was carefully aspirated and the cells were washed and lysed using 50 µL of cold ATAC-Resuspension Buffer (RSB) containing 0.1% NP40, 0.1% Tween-20 and 0.01% digitonin by pipetting up and down three times and incubating the cells for 3 minutes on ice. The lysis was washed out by adding 1 mL of cold ATAC-RSB containing 0.1% Tween-20 and inverting the tube three times. Nuclei were pelleted at 500 RCF for 10 minutes at 4°C, the supernatant was carefully removed and nuclei were resuspended in 50 µL of transposition mixture (25 µL 2x TD buffer, 2.5 µL transposase (100 nM), 16.5 µL DPBS, 0.5 µL 1% digitonin, 0.5 µL 10% Tween-20, 5 µL H2O) by pipetting six times up and down, followed by 30 minutes incubation at 37°C at 1,000 RPM mixing rate. After MinElute clean-up and elution in 21 µL elution buffer, the transposed fragments were pre-amplified with Nextera primers by mixing 20 µL of transposed sample, 2.5 µL of both forward and reverse primers (25 µM) and 25 µL of 2x NEBNext Master Mix (program: 72°C for 5 minutes, 98°C for 30 seconds and 5 cycles of [98°C for 10 seconds, 63 °C for 30 seconds, 72°C for 1 minute] and hold at 4°C). To determine the required number of additional PCR cycles, a qPCR was performed. The final amplification was done with the additional number of cycles, samples were cleaned-up by MinElute and libraries were finalized using the KAPA Library Quantification Kit as previously described. Samples were sequenced on a NextSeq500, except the MM099 baseline sample which was sequenced on NovaSeq6000.

#### Sequencing data processing

ATAC-seq reads were mapped to the reference genome (hg19) using Bowtie2 (v2.2.6) (Langmead and Salzberg 2012), post-processed with Picard (v1.134) (Broad Institute 2019) to remove duplicate reads, and SAMtools (v1.8) (H. Li et al. 2009) to retain reads with high mapping quality (Q30). BigWig files for visualization in UCSC and IGV were created with deepTools2 (Ramírez et al. 2016). Peaks were called using the MACS2 algorithm (q < 0.05, --nomodel) (Zhang et al. 2008).

#### Data analysis

ATAC-seq peaks were filtered as described in (Corces et al. 2018). Briefly, summit calls of each sample were extended 250 base-pairs in each direction to obtain fixed-width peaks of 501 base-pairs. Overlapping fixed-width peaks within each sample were filtered iteratively using the peak scores (peaks were ranked by the peak score, and any overlapping peak was filtered, then the same is done for the next high-scoring peak and so on). After obtaining sets of non-overlapping fixed-width peaks per sample, peak scores within each sample were normalized by the number of fragments in peaks. Next, the peaks of nine cultures (together with the peaks from eight SOX10-KD experiments) were combined and filtered iteratively using the process described above with normalized peak scores. This process resulted in 355,951 consolidated peaks. ATAC-seq signal was quantified across the consolidated peak set using the featureCounts commands (Liao, Smyth, and Shi 2014). ATAC-seq counts over consolidated peak set were further processed in R/Bioconductor (library-size normalized using edgeR (McCarthy, Chen, and Smyth 2012) and quantile-normalized using the preprocessCore package (Bolstad [2014] 2018)). In total, 144,017 differentially accessible peaks were obtained after performing a likelihood-ratio test using DESeq2 (Love, Huber, and Anders 2014b) with FDR cut-off 0.05. Hierarchical clustering on the differentially accessible peaks was performed using fastcluster (Müllner 2013) and heatmaps were generated using pheatmap (Kolde 2019). Genomic annotation of the peaks was done with the ChIPseeker package (Yu, Wang, and He 2015). Motif content of each cluster was determined using HOMER (Heinz et al. 2010) with the full set of 355,951 consolidated peaks as background.

### Public data

#### Analysis of publicly available ATAC-seq data

Normalized ATAC-seq signal from 410 samples across 562709 regulatory regions were downloaded from (Corces et al. 2018). Regulons of EGR3, RXRG and NFATC2 were analyzed with i-cisTarget (Imrichová et al. 2015) to obtain candidate regulatory regions. The predicted regions were then overlapped with the set of consolidated peaks and ATAC-seq signal visualized as violin plots in Figure S5. Invasive and proliferative regions predicted in Verfaillie et al were processed the same way and visualized in Figure S5 as well.

#### Analysis of publicly available expression data

The normalized expression matrix for single-cell RNA-seq data of melanoma biopsies was downloaded from GEO (GSE115978). We used gene rankings of each cell and calculated AUC values for regulons identified in the analysis of ten baselines (Aibar et al. 2017) (**Supplementary File 1**). The dataset contains malignant cells from 33 patient biopsies (29 metastasis and 4 primary tumors). Bulk RNA-seq data of melanoma samples were analyzed as described previously (Verfaillie et al. 2015). Similar to the analysis of scRNA-Seq data, gene rankings per sample were used for calculating AUC values for baseline regulons.

#### Analysis of publicly available ChIP-seq data

Raw reads for publicly-available ChIP-seq datasets (listed in the table below) were downloaded from SRA using SRA toolkit. Reads were mapped to reference genome using Bowtie2, post-processed with Picard to remove duplicates and SAMtools to remove reads with mapping quality below 30. Bigwig files were created using Deeptools (with cpm normalization). Corresponding input or whole cell extract reads were also downloaded for peak calling, and MACS2 was used for peak calling (with p-value 0.01 and q-value 0.05 thresholds). Aggregation plots and heatmaps of ChIP-seq signal were generated using Deeptools.

#### Public data sets used in this study

The following publicly available ChIP-seq data were used in this study.

| **Accession Number** | **Description** |
| --- | --- |
| GSM3144740-3144747 | OmniATAC-seq in MM057 and MM087 at multiple time points (0, 24, 48 and 72 hours) after knockdown of SOX10 |
| GSM1517752 | SOX10 ChIP-seq in 501Mel cell line |
| GSM803500 | JUND ChIP-seq in HEPG2 cell line |
| GSM659790 | FOS ChIP-seq in MCF7 cell line |
| GSE115978 | scRNA-seq of melanoma patients |
| The raw count matrix composed of 375 samples was downloaded from the GDAC Firehose data portal (stddata, timestamp: 16_03_2014, cohort: SKCM). | Bulk RNA-seq of melanoma patients |
| Normalized ATAC-seq signal from 410 samples across 562709 regulatory regions was downloaded as a matrix from the UCSC Xena data portal. | Bulk ATAC-seq of melanoma patients |

## Data and Software Availability

A SCope instance containing this data is available online at http://scope.aertslab.org/#/Wouters_Human_Melanoma. A UCSC hub with bigwig and BED files of our ATAC-seq data is available via: http://ucsctracks.aertslab.org/papers/wouters_human_melanoma/hub.txt. The scRNA-seq and ATAC-seq data has also been deposited in GEO under accession number GSE134432. Raw images and tracking information for the single-cell migration experiments are made publicly available on the Open Science Framework (OSF) of the Center for Open Science (COS) at http://osf.io (DOI 10.17605/OSF.IO/E6AHM).

## Code Availability

No custom software was used for the analyses in this study. All publicly available tools and packages (and their versions and parameters used) are mentioned in the Methods.

## List of supplementary materials

**Supplementary Notes**

Supplementary Note 1. Genetic demultiplexing of scRNA-seq data using demuxlet

**Supplementary Files**

Supplementary File 1. All regulons as identified by SCENIC (SCENIC_regulons.gmt)

**Supplementary Tables**

Supplementary Table S1. Summary statistics of cell quality filtering.

Supplementary Table S2. Sequencing details and summary statistics of single-cell sequencing runs.

Supplementary Table S3. Custom primers used for library preparation and sequencing.

## Supplementary Figures

**Supplementary Figure S1**. Functional characterization of ten patient-derived melanoma cultures. **a**, The heatmap shows the activity of literature-derived gene signatures (rows) in each cell (columns), measured by AUCell. Unsupervised hierarchical clustering demonstrates that two groups are formed based on contrasting activity of melanocyte and pigmentation related signatures vs. de-differentiated (mesenchymal-like, neural crest-like, immune-like) and resistance-related signatures. **b**, Each melanoma culture has a subpopulation with high cell cycle activity, shown in a t-SNE plot of the 10 melanoma cultures colored according to G2M checkpoint gene signature activity from the hallmark collection (MSigDB) (Liberzon et al. 2015). **c**, GSEA for the first two principal components demonstrates that PC1 separates cells with high melanocyte-like activity from cells with high mesenchymal-like activity, while PC2 correlates with a gradient of immune-related processes. **d**, GSEA for the first two canonical correlation dimensions demonstrates that CC1 correlates, similar to PC1, with a melanocyte-like to mesenchymal-like gradient, while CC2 is associated with cell cycle activity.

**Supplementary Figure S2.** The Gini coefficient identifies genes with heterogeneous expression within each melanoma culture. **a**, The normalized Gini coefficient for each gene in each MM line is plotted and genes that are present in the mesenchymal-like and the melanocyte-like gene signatures from Widmer et al. 2012 are indicated in green and red, respectively. GSEA results in significant enrichment of the melanocyte-like signature in the mesenchymal-like cell lines (with FDR<0.05). **b**, The two scRNA-seq methods 10x and Drop-seq yield comparable results on the cell line MM057, as shown by a CCA plot. **c**, The Gini coefficients of 10x and Drop-seq on MM057 are correlated (pearson r = 0.38).

**Supplementary Figure S3.** The transcriptional activity of a cell migration signature predicts the migratory capacity of melanoma cultures. Same as **Figure 2c** with addition of MM074 lines (center line, median; box limits, upper and lower quartiles).

**Supplementary Figure S4.** Functional characterization of three cell states and associated regulons using additional data. **a**, Violin plot for MITF regulon activity is shown for 10 melanoma cultures demonstrating a gradual decrease from fully melanocyte-like cultures (MM001, MM011, MM031) to intermediate cultures (MM074, MM087, MM057). **b**, Normalized ATAC-seq signal in melanocyte-like regions (n=6669) and mesenchymal-like regions (n=13453) as previously identified by Verfaillie et al show higher chromatin accessibility in melanocyte like regions in melanocyte-like and intermediate cultures while lower chromatin accessibility in mesenchymal-like regions, and vice versa. **c**, Normalized ATAC-seq signal in SOX10 bound, FOS bound and JUND bound regions, as previously identified by ChIP-seq (REFs) show contrasting chromatin accessibility between mesenchymal-like cultures vs melanocyte-like for SOX10 bound regions, while a gradual decreased accessibility in AP-1-associated regions (FOS and JUND) going from mesenchymal-like, to intermediate and finally to melanocyte-like cultures. **d**, Normalized ATAC-seq signal is shown at IRF2 (top left), FN1 (top right), SOX9 (bottom left) and NFATC2 (bottom right) gene loci show higher accessibility in intermediate and mesenchymal-like cultures compared to the melanoctye-like cultures. **e**, Boxplot for transcriptional activity of NFATC2, EGR3 and RXRG regulons in melanoma biopsies (Jerby-Arnon et al. 2018) seperated and colored according to origin of resection (metastasis or primary lines; center line, median; box limits, upper and lower quartiles). **f**, Scatter plot of transcriptional activity between NFATC2, RXRG and EGR3 regulons (on the top row) indicate a positive correlation in regulon activity (sizes of regulons are 40 (NFATC2), 103 (EGR3) and 35 (RXRG) with the overlap between regulons being 10 (NFATC2 & EGR3), 11 (EGR3 & RXRG) and 2 (NFATC2 & RXRG)). Scatter plot of transcriptional activity between HES6 and NFATC2, EGR3, and RXRG regulons show no correlation between these regulons. **g**, Scatter plots for regulons same as (**f**) but with using bulk RNA-seq data from TCGA-SKCM set indicate again a positive correlation between NFATC2, RXRG and EGR3 regulons on the top row with correlation coefficients of 0.55, 0.54 and 0.60 (significant at p-value threshold of 0.05). Bottom row shows scatter plots for regulon activity for HES6 versus NFATC2, EGR3 and RXRG with weak but significant negative correlation (−0.38, −0.21 and −0.22). Cells are colored according to the classification from Verfaillie et al. (INV is for invasive or mesenchymal-like, PRO is proliferative or melanocyte-like, and TIL is for samples high T-cell infiltration)

**Supplementary Figure S5.**

Violin plots for melanocyte-like, mesenchymal-like, NFATC2-linked, RXRG-linked, EGR3-linked and intermediate cell state-linked enhancers (union of NFATC2, RXRG and EGR3 linked enhancers) were generated using ATAC-seq data of TCGA. ACC: adrenocortical carcinoma; BLCA: bladder urothelial carcinoma; BRCA: breast invasive carcinoma; CESC: cervical squamous cell carcinoma; CHOL: cholangiocarcinoma; COAD: colon adenocarcinoma; ESCA: esophageal carcinoma; GBM: glioblastoma multiforme; HNSC: head and neck squamous cell carcinoma; KIRC: kidney renal clear cell carcinoma; KIRP: kidney renal papillary cell carcinoma; LGG: low grade glioma; LIHC: liver hepatocellular carcinoma; LUAD: lung adenocarcinoma; LUSC: lung squamous cell carcinoma; MESO: mesothelioma; PCPG: pheochromocytoma and paraganglioma; PRAD: prostate adenocarcinoma; SKCM: skin cutaneous melanoma; STAD: stomach adenocarcinoma; TGCT: testicular germ cell tumors; THCA: thyroid carcinoma; UCEC: uterine corpus endometrial carcinoma.

**Supplementary Figure S6.** Comparative alignment of transition trajectories across different methods. Trajectories predicted by DiffusionMap, Scorpius and Monocle were aligned by applying dynamic time warping cellAlign (Alpert et al. 2018) and the predicted optimal alignment is shown with a white line. The concordance between different methods is high (with the exception of MM057)

**Supplementary Figure S7**. Recurrent transition trajectory across melanoma cultures. **a**, Interpolated and scaled gene expression (see Methods) after SOX10-KD for each culture along pseudo time shows a collapse of the transcriptome (6,495, 6,396 and 6,113 in MM057, MM074 and MM087, respectively), and up-regulated expression of 1,209, 1,308 and 1,591 genes, respectively. **b-c**, Ternary plots (see Methods) for gene expression (**b**) and gene signature activity (**c**) after SOX10-KD indicating very high transcriptional concordance between melanoma cultures and various relevant down- and up-regulated processes. **d**, Heatmap with the activity of literature-derived gene signatures (rows) in each cell (columns), measured by AUCell for all melanoma cultures after SOX10-KD (for both 10x and Drop-seq scRNA-seq technologies), indicating the recurrent down-regulation of cell cycle and melanocytic transcriptional programs, and up-regulation of cellular migration, the epithelial-to-mesenchymal transition, cancer metastasis, immune cell activation, angiogenesis, and melanoma-specific gene sets such as the signatures for the SOX9-positive cultures (Hoek et al. 2006; Widmer et al. 2012, Verfaillie et al. 2015), the melanoma TNF response (Riesenberg et al. 2015), the AXL program signature (Tirosh et al. 2016) and the signature for acquired resistance to BRAF inhibition (Shaffer et al. 2017). Comparison between 10x and Drop-seq scRNA-seq modalities demonstrates the consistency of the observed transcriptional changes.

**Supplementary Figure S8.** Heatmap showing regulon activity in MM074 for regulons that are common (i.e. assigned to the same TF) between MM057, MM074 and MM087 (n=114). Cells are ranked according to diffusion maps’ pseudotime and regulons are clustered using unsupervised hierarchical clustering.

**Supplementary Figure S9.** Heatmap showing regulon activity in MM087 for regulons that are common (i.e. assigned to the same TF) between MM057, MM074 and MM087 (n=114). Cells are ranked according to diffusion maps’ pseudotime and regulons are clustered using unsupervised hierarchical clustering.

**Supplementary Figure S10.** Heatmap showing regulon activity in MM057 for regulons that are common (i.e. assigned to the same TF) between MM057, MM074 and MM087 (n=114). Cells are ranked according to diffusion maps’ pseudotime and regulons are clustered using unsupervised hierarchical clustering.

**Supplementary Figure S11.** Gene regulatory networks (GRNs) as identified by SCENIC in MM074 (top), MM087 (middle), and MM057 (bottom) colored by expression changes over time (from baseline to non-template control to SOX10-KD at 24, 28 and 72 hours).

**Supplementary Figure S12.** Heatmap for 114 genes that are reported to be downregulated after THZ1 (Eliades et al. 2018) also show downregulation after THZ2 (compared to DMSO treatment).

## Acknowledgements

This work was funded by an ERC Consolidator Grant to S. Aerts (no. 724226_cis-CONTROL), and by the KU Leuven (grant no. C14/18/092 to S. Aerts), the Harry J. Lloyd Charitable Trust, the Foundation Against Cancer (grant no, 2016-070 to S. Aerts), PhD fellowships from the FWO (L.M., no. 1S03317N) and a postdoctoral research fellowship from Kom op tegen Kanker (Stand up to Cancer), the Flemish Cancer Society (J.W). Computing was performed at the Vlaams Supercomputer Center. Single-cell infrastructure was funded by the Hercules Foundation (grant no. AKUL/13/41). The funders had no role in study design, data collection and analysis, decision to publish or preparation of the manuscript.

## Author Contributions

J.W., Z.K.A. and S. A. conceived the study. J.W. performed the experimental work for the 10x scRNA-seq datasets with the help of V.C., S.M. and S.P.. J.W. performed the experimental work for the Drop-seq datasets with the help of K.D. and V.C.. J.W. and L.M. performed the experimental work for the OmniATAC datasets with the help of D.M. and V.C.. L.M. performed the experimental work and analysis of the single-cell migration assays with the help of F.C. and S.P.. Z.K.A, J.W. and K.I.S. performed all bioinformatics analyses with the help of G.H., M.D.W., K.D. and C.B.G.B.. A.A. and G.G. provided valuable melanoma cultures. M.D., F.R. and J.C.M. provided valuable materials and contributed to the manuscript. J.W., Z.K.A., K.I.S. and S.A. wrote the manuscript.

