## Supplementary Figures for "Single-cell gene regulatory network analysis reveals new melanoma cell states and transition trajectories during phenotype switching"

### Supplementary Figure S1

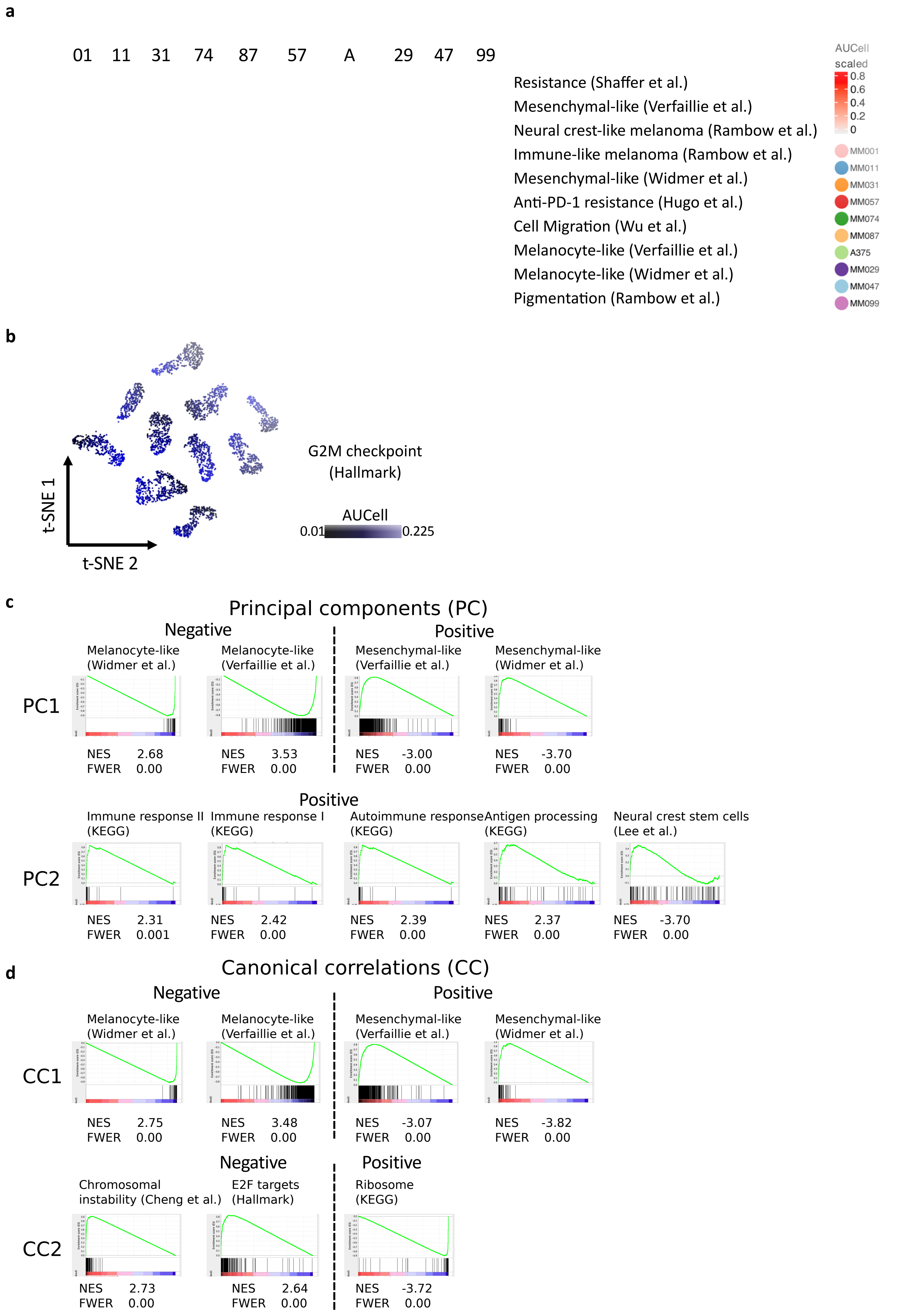

### Supplementary Figure S2

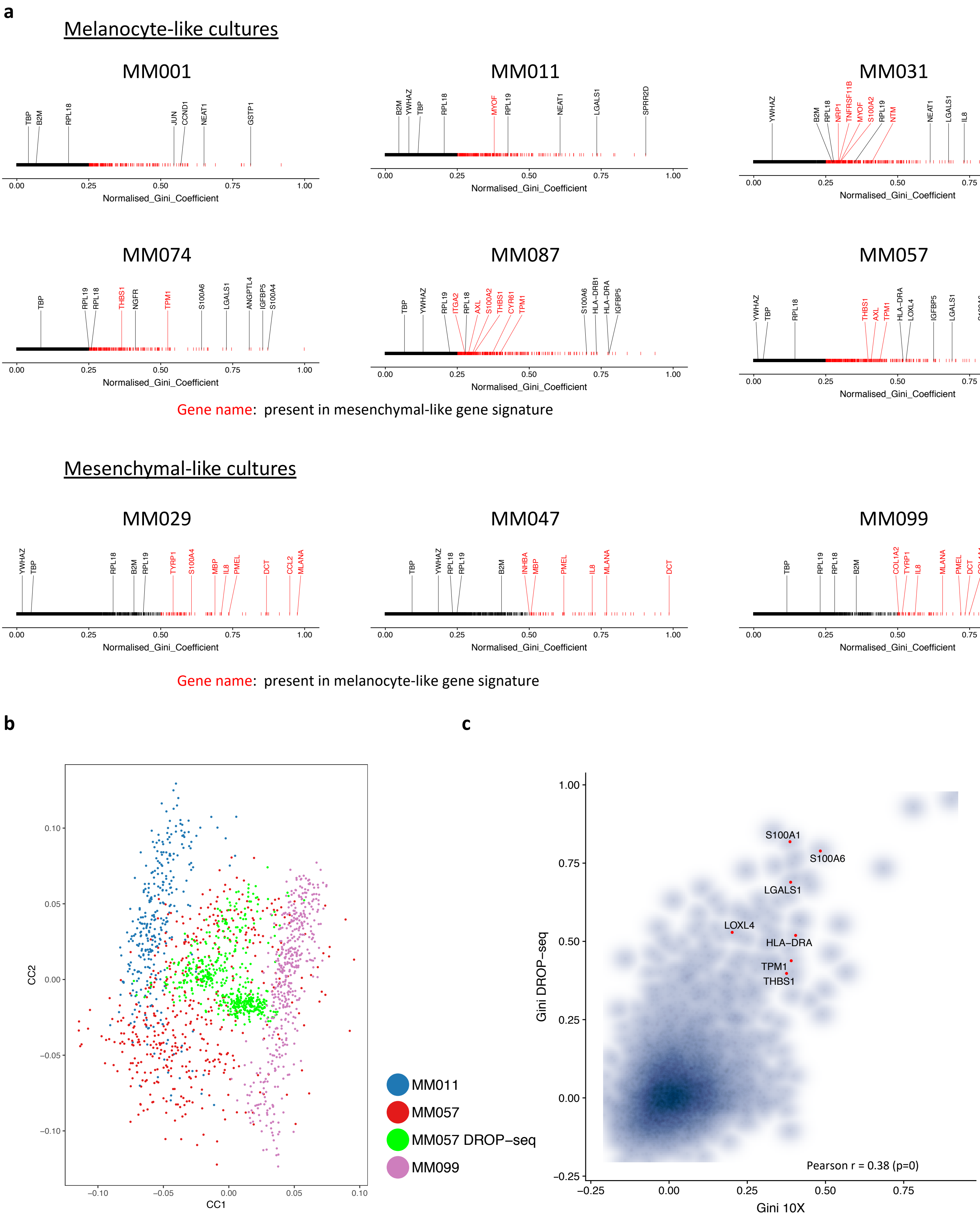

**Supplementary Figure S3**

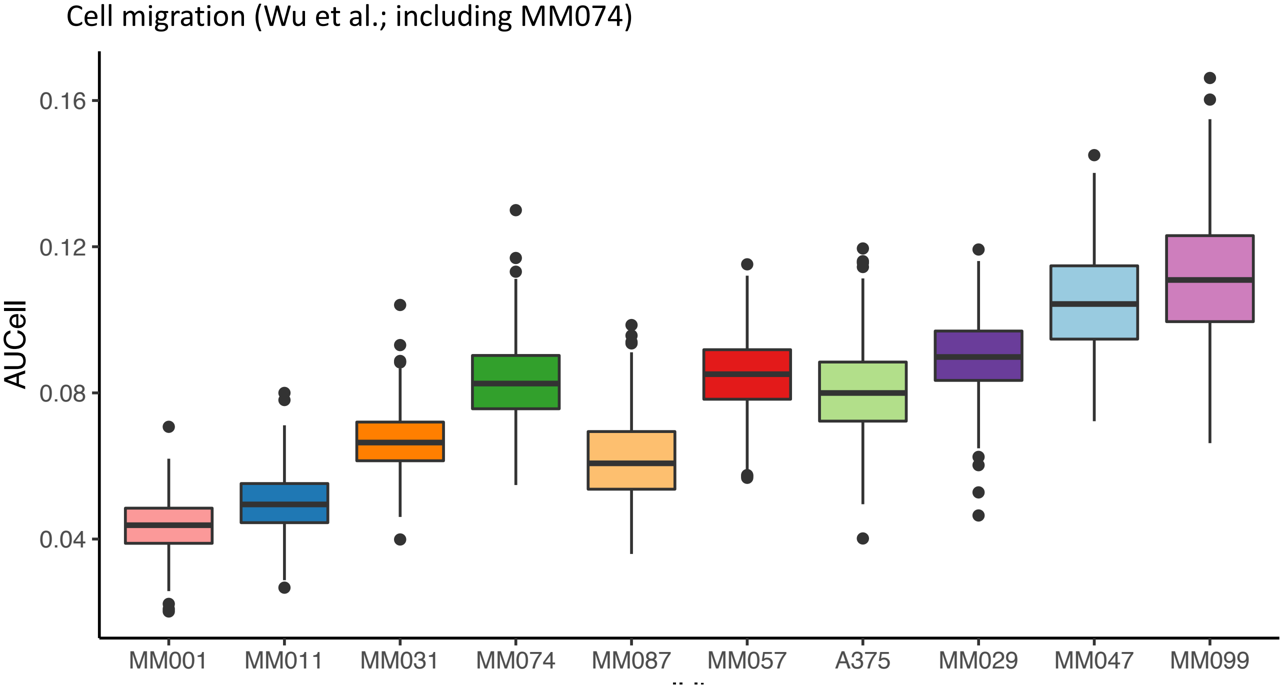

#### Supplementary Figure S4

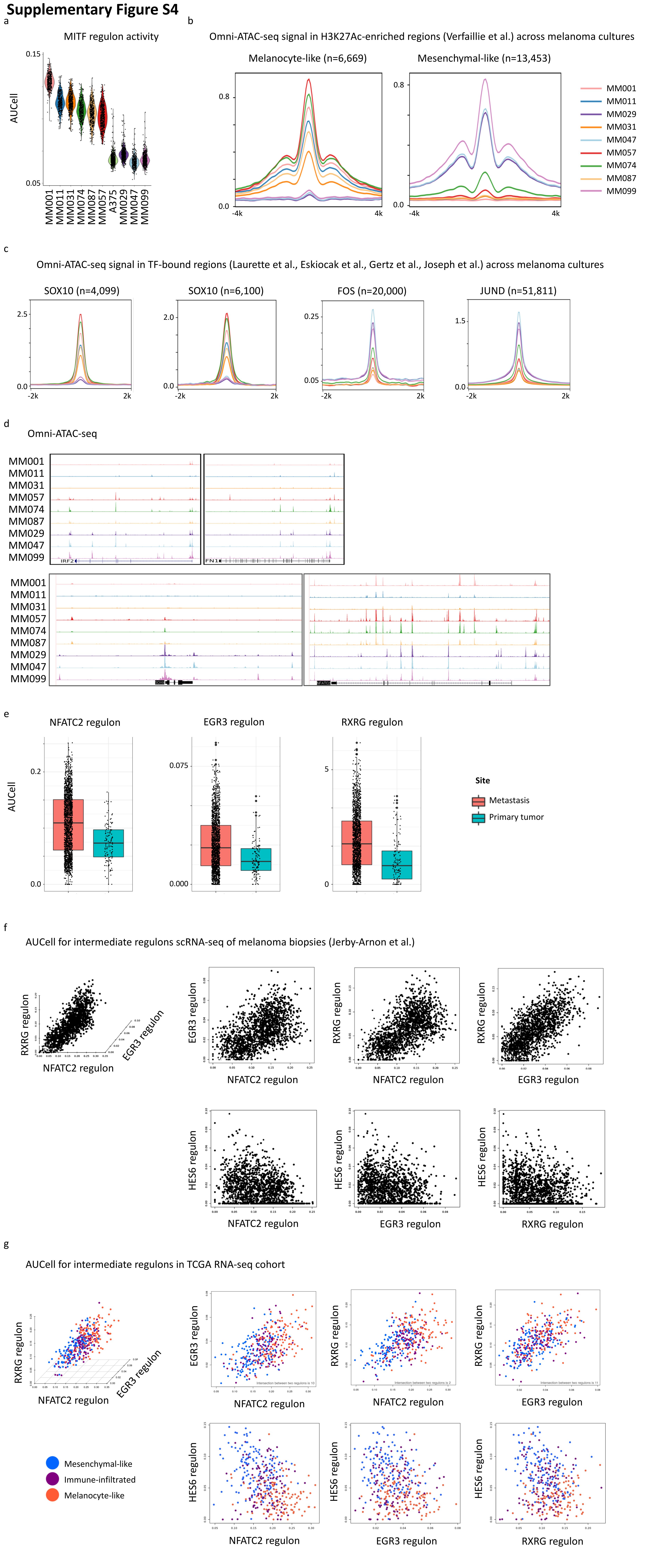

### Supplementary Figure S5

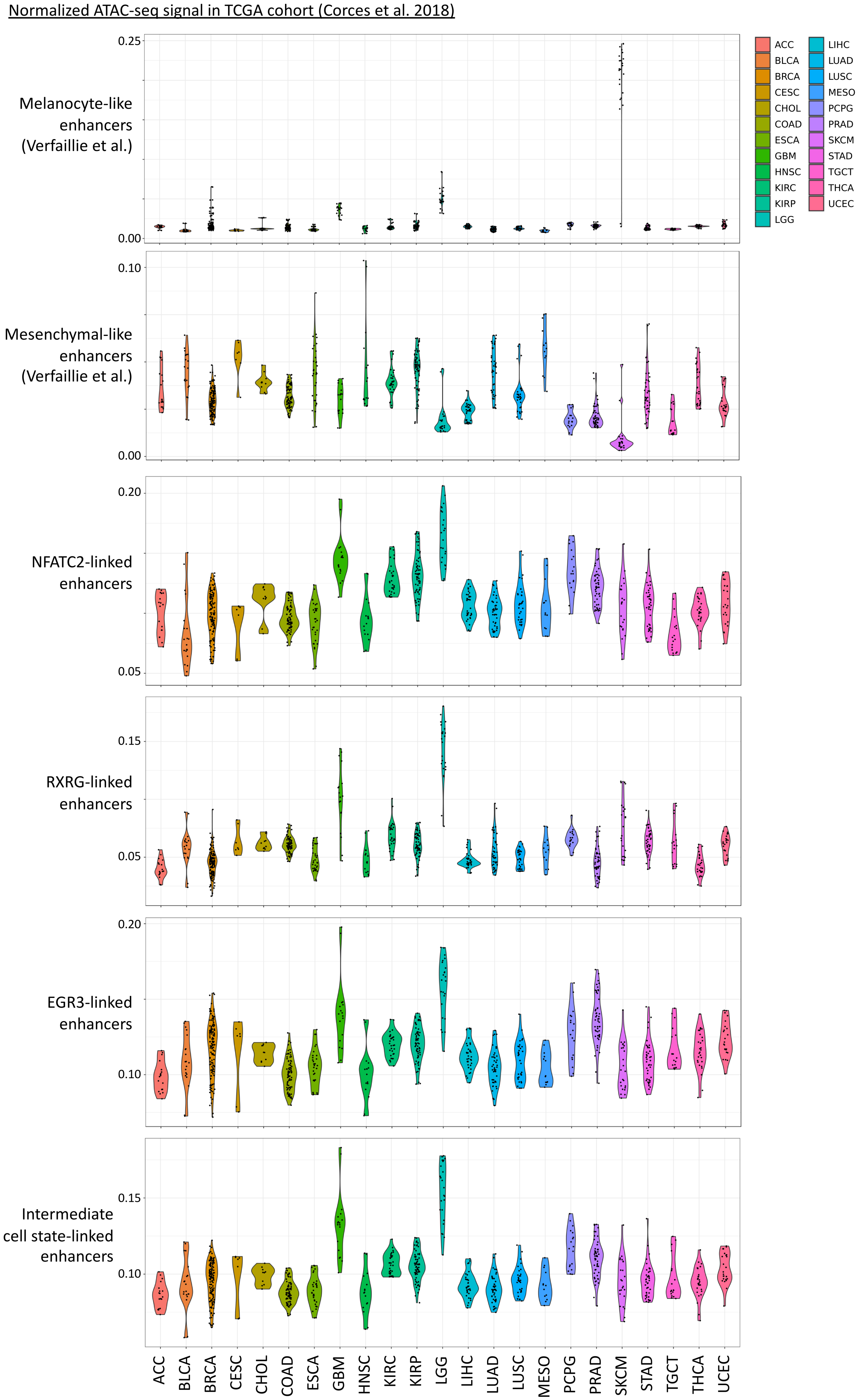

### Supplementary Figure S6

Alignment of trajectory methods

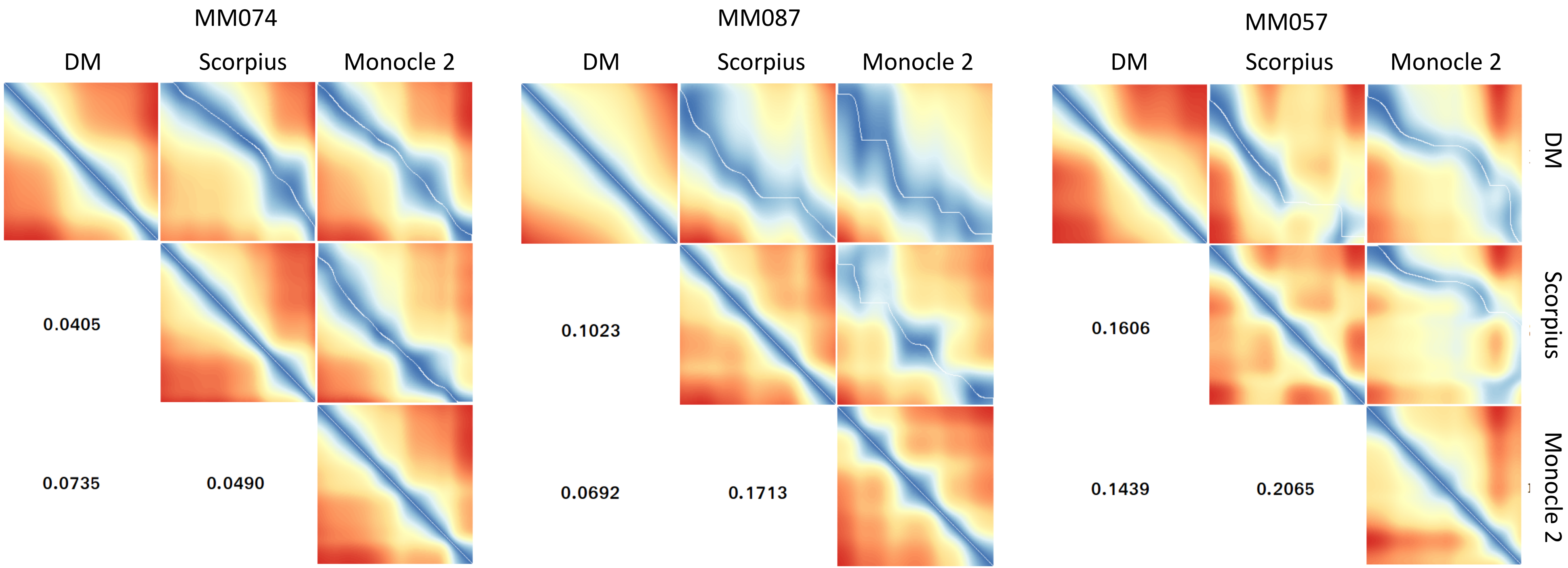

### Supplementary Figure S7

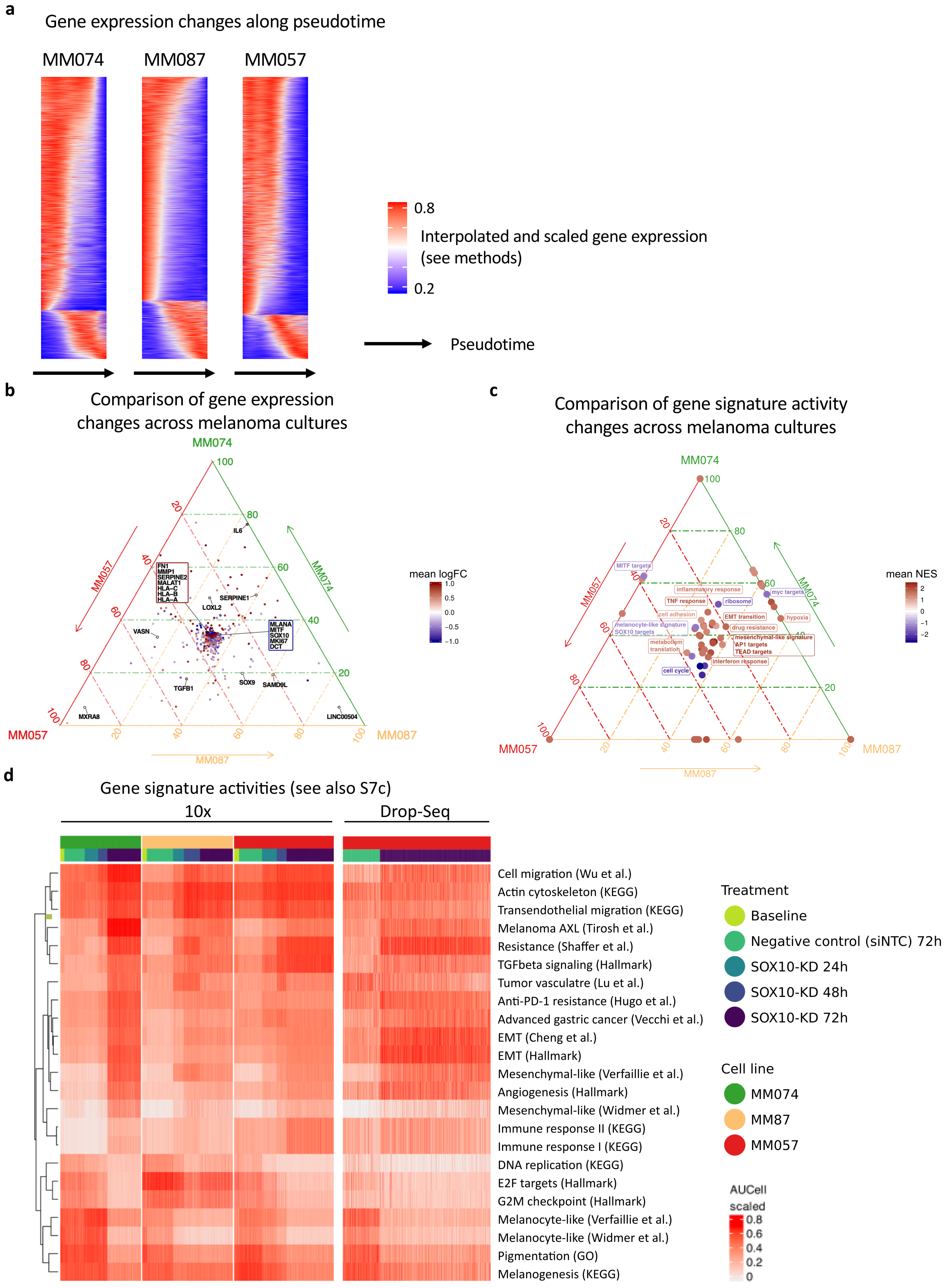

### Supplementary Figure S8

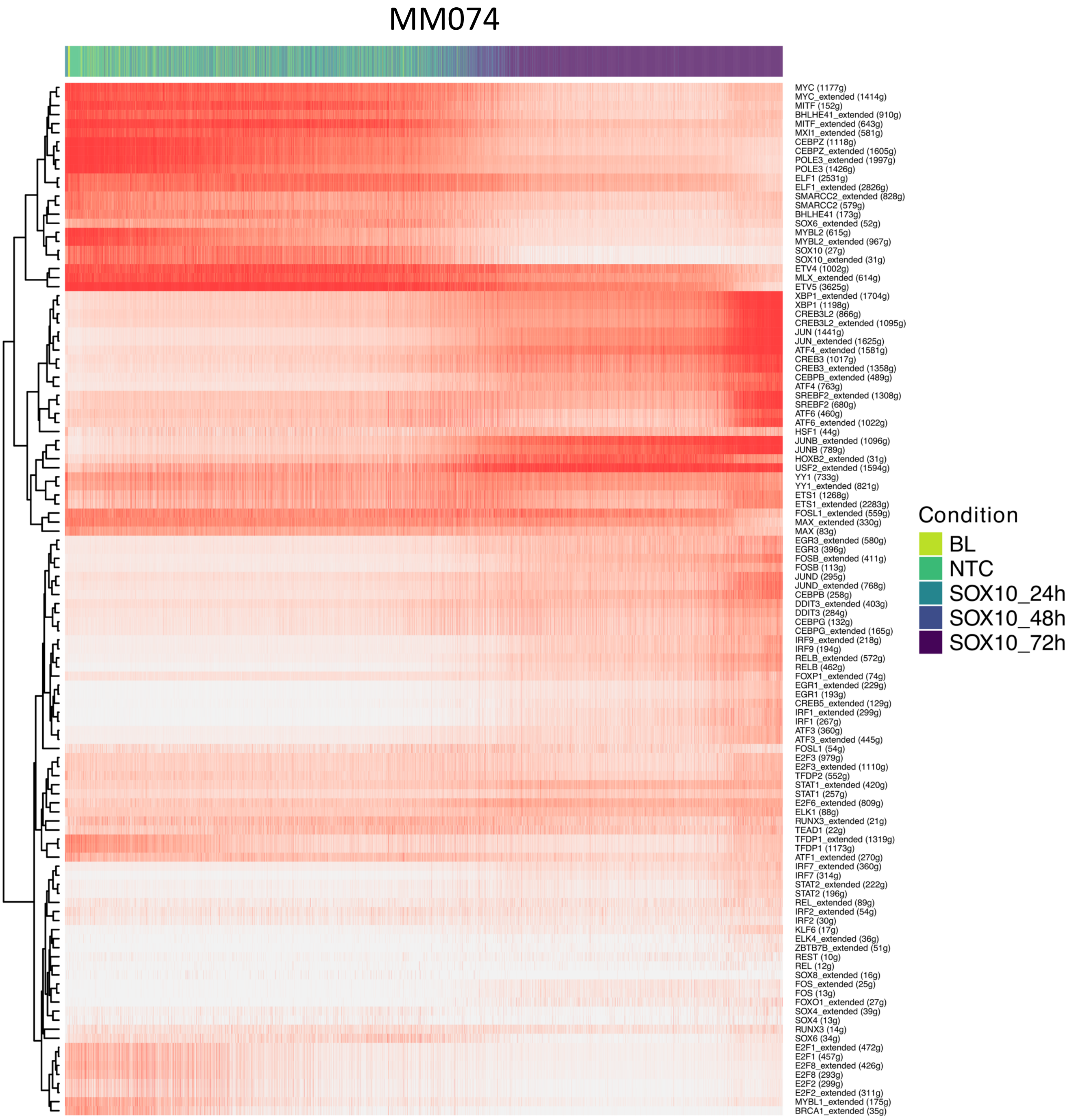

### Supplementary Figure S9

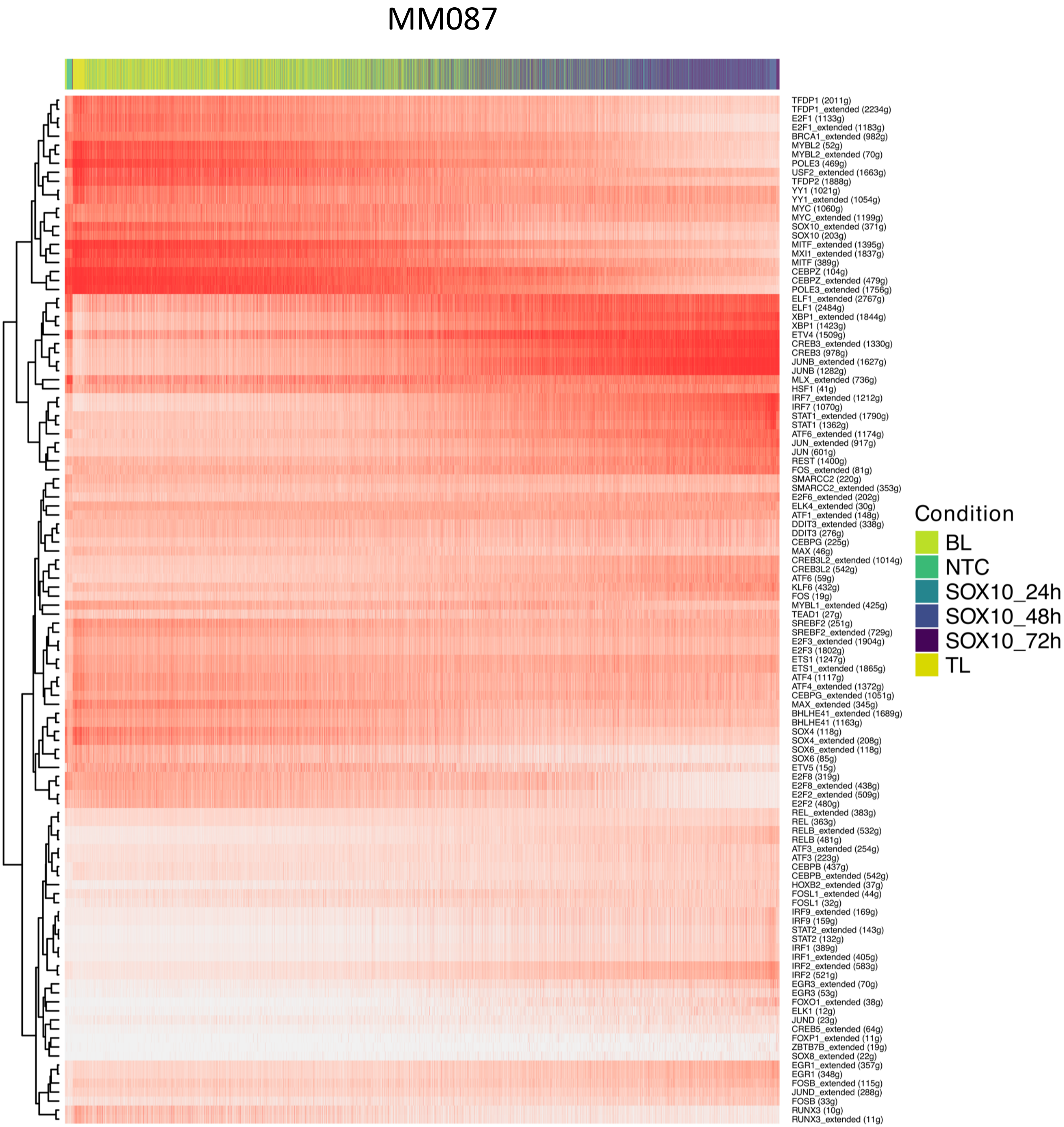

### Supplementary Figure S10

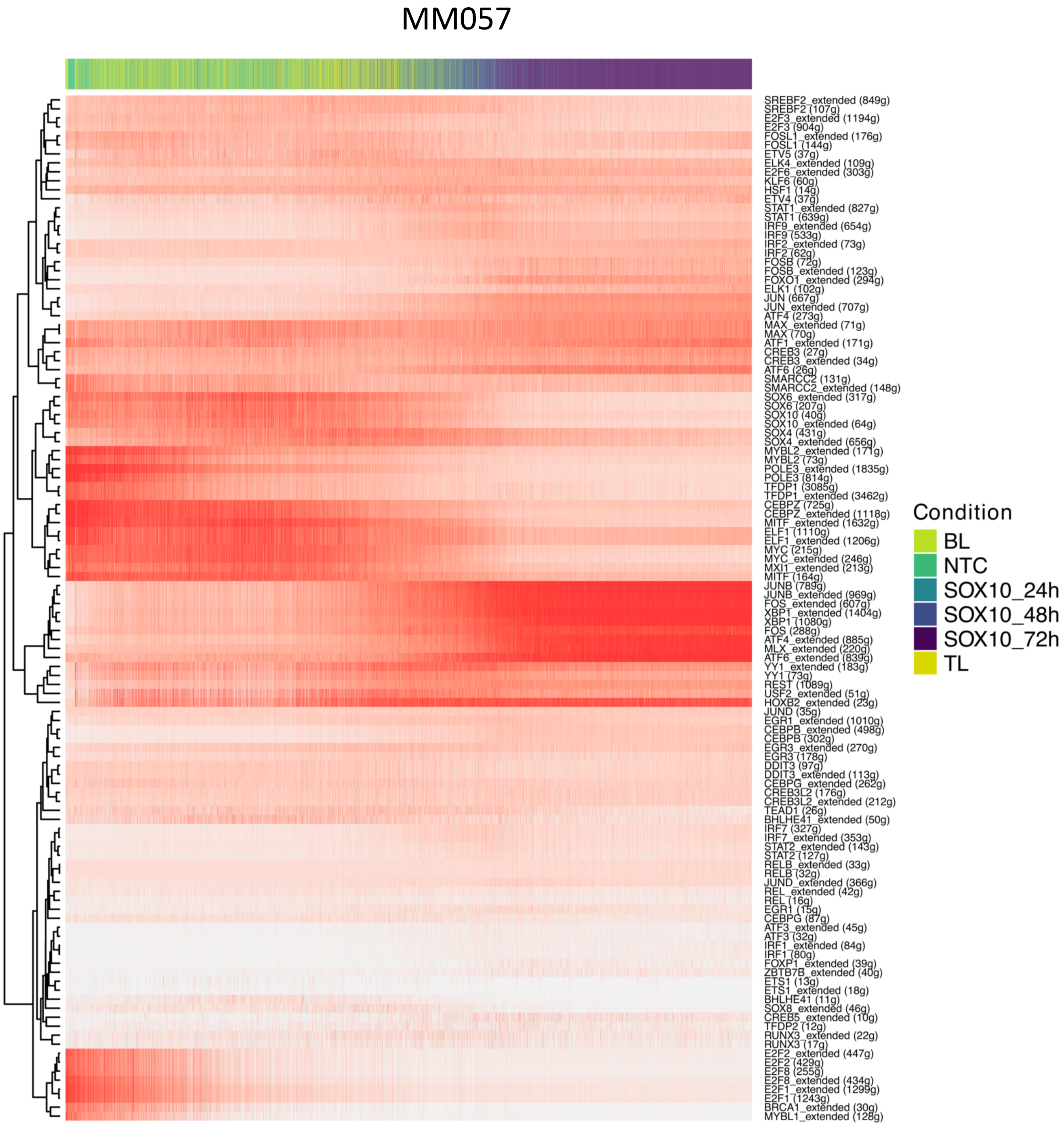

**Supplementary Figure S11**

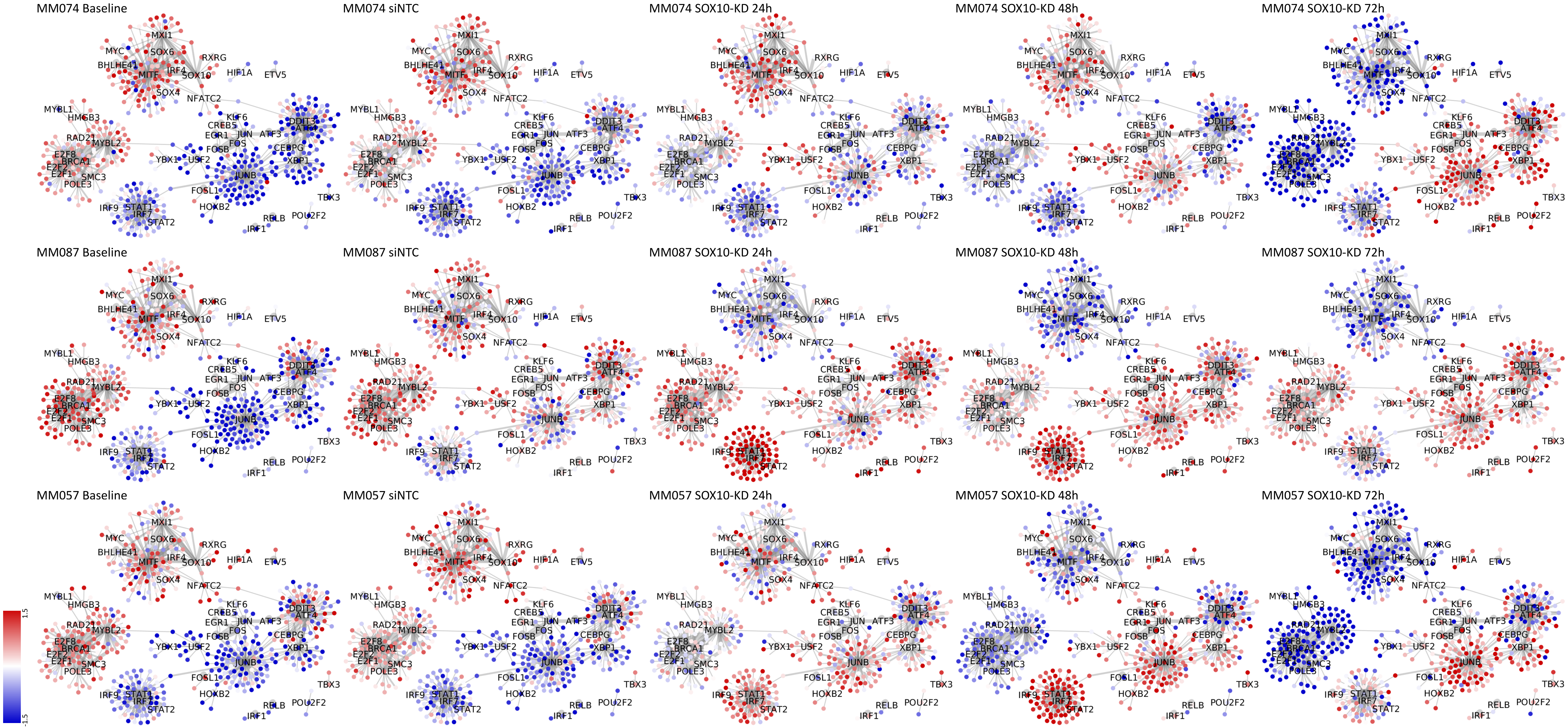

### Supplementary Figure S12

#### Comparison to public data

114 genes downregulated after 6h of THZ1 treatment

(Eliades et al., JID, 2018)

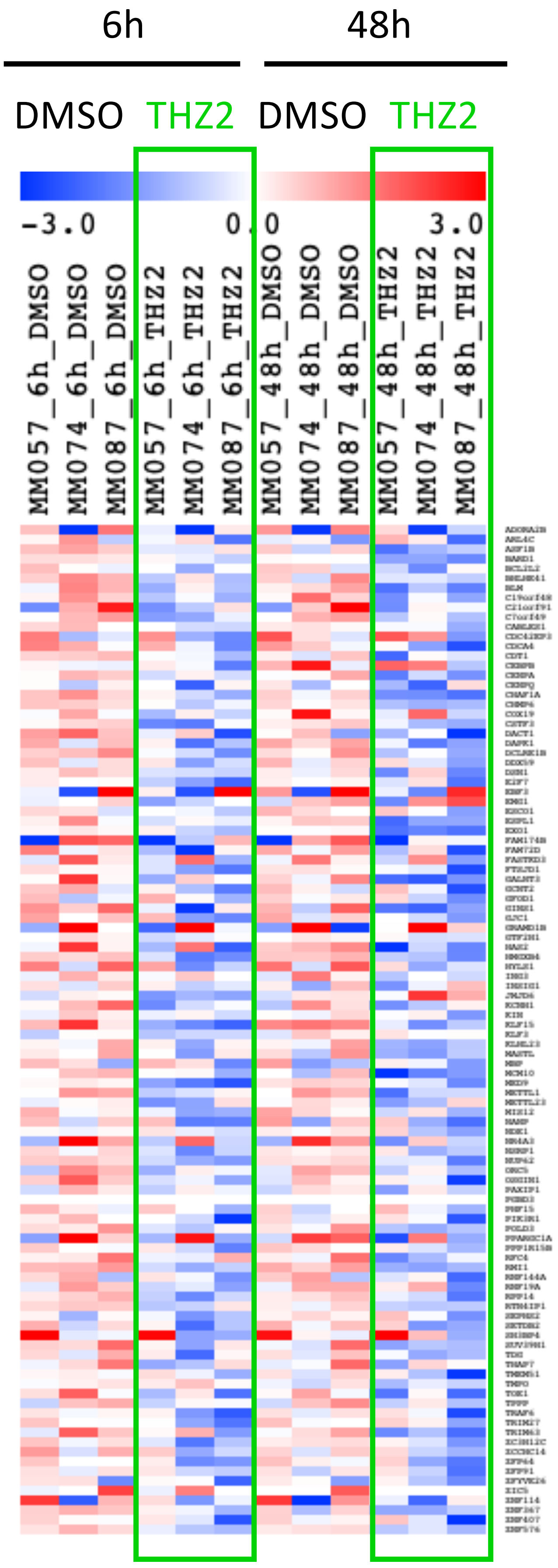
