## Supplementary Note 1 for "Single-cell gene regulatory network analysis reveals new melanoma cell states and transition trajectories during phenotype switching"

**Supplementary Note 1.** Genetic demultiplexing of single-cell RNA-seq data using demuxlet

In four of the 13 experiments we performed, we pooled melanoma cultures into a single 10x Chromium lane (**Supplementary Table 1)**. To determine the identity of the cells in these runs, we used sample-specific genetic mutations and demuxlet (Kang et al. 2018). Demuxlet predicts the likelihood of observing a set of sample-specific SNPs in each cell given the set of uncorrelated SNPs with genotype probabilities. We used high-coverage bulk RNA-seq data on the same melanoma cultures to identify sample-specific SNPs. Genotype calling was performed on ten cultures with the samtools mpileup command (version 1.3.1 with options -q 1 -Q 13 -d 8000) on the bulk RNA-seq reads that were mapped and post-processed as described previously (Verfaillie et al. 2015). We filtered the resulting vcf file based on the genotype fields to obtain an uncorrelated set of SNPs.

First, to assess the accuracy of SNP-based demultiplexing, we combined the bam files of three single-sample scRNA-seq runs (72h after SOX10-KD in MM057, MM074 and MM087) to obtain a pseudo-pooled scRNA-seq run. We analysed the resulting bam file with demuxlet and compared the predictions based on sample-specific SNPs with known cell identities. demuxlet classified 13543 of the 13545 cells correctly (**Figure 1**).

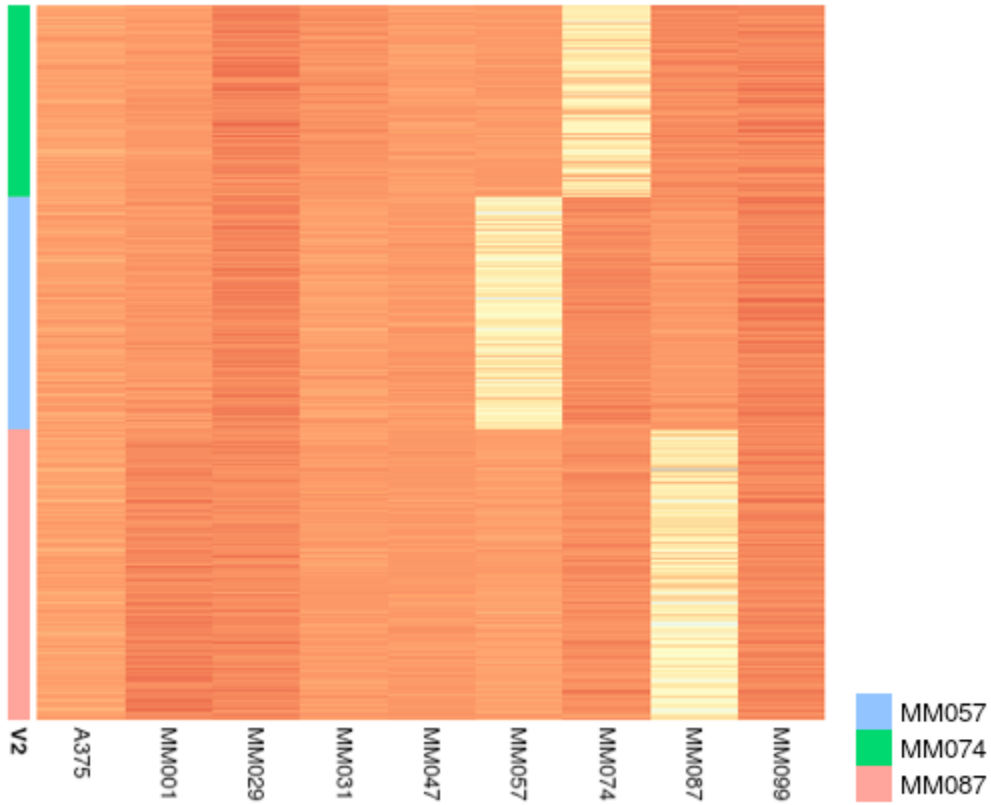


**Figure 1.** Heatmap showing log-likelihood ratios (row median-scaled) of cell identity predictions from demuxlet. On the left hand side of the heatmap, true sample identities are indicated with colored blocks (green for MM074, blue for MM057 and red for MM087).

Next, we used demuxlet on the four pooled experiments (with --alpha 0 --alpha 0.5 option to include doublet predictions as well) (**Table 1**) . Predicted doublet percentages ranged from two to five percent, with the pooled sample of ten cultures having the highest doublet percentage (5.2%). Predicted singlets were used in the downstream analyses.

| Sequencing Run | Number of demultiplexed cells | Sample Ratios | Number of predicted doublets |
|---|---|---|---|
| Baselines (10 mixed) | 4,334 | 566(A375):428(MM001):319(MM011):457(MM029):461(MM031):322(MM047):478(MM057):385(MM074):453(MM087):465(MM099) | 240 |
| 48h after SOX10-KD (3 mixed) | 3,364 | 932(MM057):901(MM074):1531(MM087) | 82 |
| 24h after SOX10-KD (3 mixed) | 3,699 | 1398 (MM057):1280 (MM074):1021(MM087) | 89 |
| 72h after NTC-KD (MM074-MM057 mix) | 4,072 | 2161(MM057):1910(MM074) | 84 |

**Table 1.** Demuxlet statistics
